# Single-molecule orientation localization microscopy for resolving structural heterogeneities between amyloid fibrils

**DOI:** 10.1101/2020.01.22.916122

**Authors:** Tianben Ding, Tingting Wu, Hesam Mazidi, Oumeng Zhang, Matthew D. Lew

## Abstract

Simultaneous measurements of single-molecule positions and orientations provide critical insight into a variety of biological and chemical processes. Various engineered point spread functions (PSFs) have been introduced for measuring the orientation and rotational diffusion of dipole-like emitters, but the widely used Cramér-Rao bound (CRB) only evaluates performance for one specific orientation at a time. Here, we report a performance metric, termed variance upper bound (VUB), that yields a global maximum CRB for all possible molecular orientations, thereby enabling the measurement performance of any PSF to be computed efficiently (~1000× faster than calculating average CRB). Our VUB reveals that the simple polarized standard PSF provides robust and precise orientation measurements if emitters are near a refractive index interface. Using this PSF, we measure the orientations and positions of Nile red (NR) molecules transiently bound to amyloid aggregates. Our super-resolved images reveal the main binding mode of NR on amyloid fiber surfaces, as well as structural heterogeneities along amyloid fibrillar networks, that cannot be resolved by single-molecule localization alone.

## I. INTRODUCTION

A key strength of single-molecule localization microscopy (SMLM) is its ability to measure the full distribution of the phenomena under study and avoid ensemble averaging. Going beyond standard SMLM to measure SM position and orientation simultaneously is critical for understanding a variety of nanoscale biological and chemical processes, such as the motions of molecular motors [1–3], the complex higher-order structures of DNA [4, 5], and reaction kinetics within catalytic nanoparticles [6]. To perform these measurements, molecular position and orientation must be encoded within the shape of the image produced by a microscope [7], i.e., its point-spread function (PSF). However, balancing the need to resolve various orientations unambiguously with the need to detect SMs efficiently with high signal-to-background ratio (SBR) remains a challenge that limits the adoption of Single-Molecule Orientation Localization Microscopy (SMOLM) for biological and chemical applications; existing techniques either cannot discriminate between similar types of molecular motions or cannot detect weak fluorescent emitters.

In order to design a PSF for measuring molecular orientation, one must have a figure of merit for comparison. The Cramér-Rao bound (CRB), which is the best-possible precision achievable by an unbiased estimator, has been used extensively for evaluating [8] and optimizing [9, 10] SMLM techniques. However, since CRB quantifies measurement performance locally, in the sense that it yields an expected measurement sensitivity for one specific molecular orientation, it is computationally expensive to characterize or optimize a PSF for all possible molecular orientations. We seek to develop an analytical framework for 1) quantifying the performance of any proposed method for measuring molecular orientation, 2) illuminating why a particular method performs well or poorly in a particular situation, and 3) comparing the sensitivity of various methods for measuring specific types of rotational motions.

In this paper, we derive a performance metric, termed variance upper bound (VUB), that globally bounds the CRB to quickly evaluate any optical method for measuring molecular orientation. Our VUB analysis surprisingly shows that a microscope with two polarization detection channels, exhibiting a polarized (standard) PSF, provides superior measurement precision when molecules are near a refractive index interface, especially when they lie perpendicular to the optical axis, even under low SBR. This conventional method of measuring molecular orientation has poor performance in index-matched samples unless a perturbation, such as defocus, is added to the optical system [11]. To demonstrate, we use a form of SMLM termed transient amyloid binding (TAB) [12] and a maximum likelihood estimator that promotes sparsity [13, 14] to measure the position and orientation of fluorescent molecules transiently bound to amyloid fibrils. For the first time to our knowledge, SMOLM reveals the orientation and rotational diffusion of Nile red (NR), a fluorescent molecule whose quantum yield increases when exposed to a nonpolar environment [15, 16], when bound to fibrils at the single-molecule level. We show that SMOLM probes structural heterogeneities between these amyloid aggregates that are not detectable by standard SMLM.

## II. ORIENTATION LOCALIZATION MICROSCOPY AND VARIANCE UPPER BOUND

To begin, we model a fluorescent molecule as a dipole-like emitter [17–19] with an orientational unit vector ***μ*** = [*μ_x_, μ_y_, μ_z_*]^*T*^, or equivalent angles (*φ, θ*) in spherical coordinates (Fig. S1). Assuming that a molecule’s rotational correlation time is faster than its excited state lifetime and the camera integration time [20], its orientation trajectory can be modeled by a second-moment vector 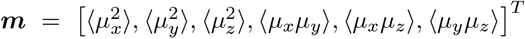, where each entry is a time-averaged second moment of ***μ*** [21, 22]. These orientational second moments contain information on both the orientation of a molecule and its “wobble” or rotational diffusion, represented by the solid angle Ω, within a camera frame. Fluorescence photons collected from such an emitter are projected to a microscope’s image plane with an *n*-pixel intensity distribution that can be modeled as a linear superposition of six basis images weighted by ***m***, i.e.,

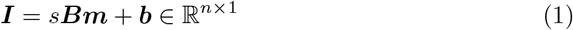

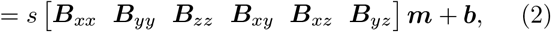

where *s* is the number of photons collected by the detector and ***b*** is the number of background photons in each pixel. Each so-called basis image ***B**_j_* ∈ ℝ^*n×*1^, corresponding to the response of the optical system to each orientational second-moment component *m_j_*, can be calculated by vectorial diffraction theory [21–24], and various optical elements, such as phase masks, polarizers, and waveplates, can be used to encode orientation information within ***B***_*j*_ more efficiently. A key feature of this forward imaging model is its linearity with respect to the orientation parameters ***m*** of interest; ***B*** is simply a linear operator or transformation from object space to image space, and the fundamental measurement sensitivity of the optical system is bounded by considering ***B*** directly.

We next quantify the Fisher information (FI) contained in the basis images, i.e., the amount of information that the basis images contain about each of the orientational second moments. A larger FI corresponds to a smaller CRB and higher sensitivity for measuring ***m***. Given our linear imaging model, the FI matrix ***J*** ∈ ℝ^6×6^ for the orientational second moments can be written as (Supplementary Material 1A)

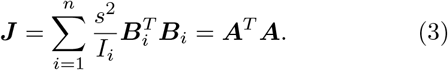

where ***B***_*i*_ ∈ ℝ^1×6^ is the *i*^th^ row of ***B***. Here, we introduce ***A*** ∈ ℝ^*n*×6^, where each entry of ***A*** can be bounded by (Supplementary Material 1A):

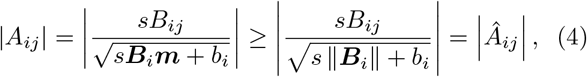

Since photon detection is a Poisson process, the entry *A_ij_* represents the square root of the expected signal-to-noise ratio of the *j*^th^ orientational second moment measured by the *i*^th^ camera pixel.

Note that ***J*** is a function of ***m*** and may vary dramatically for different molecular orientations. We now develop our VUB matrix **Γ** (Supplementary Material 1A) to bound the CRB matrix ***R*** = ***J*** ^−1^ as,

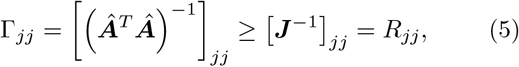

where the subscript *jj* denotes the *j*^th^ diagonal element. Since **Γ** is independent of ***m***, i.e., it is only a function of the imaging system and not molecular orientation, VUB can be used to efficiently estimate the global performance of the imaging system and the precision of measuring each orientational second moment under various SBRs without expensive calculations of CRB.

We calculated the trace Γ_Σ_ of **Γ** as a summary measure of performance of seven orientation-sensing PSFs, namely unpolarized standard, polarized standard, bifocal microscope [25], bisected [26], double helix [27], tri-spot [28], and quadrated [29], for a molecule emitting at a glass-water interface (Fig. 1(a)). To quantify the accuracy of our VUB, we compared Γ_Σ_ to the minimum/average/maximum value of the trace *R*_Σ_ of all diagonal entries of the CRB matrix over all of orientation space (Fig. 1(a) inset). We find Γ_Σ_ estimates the average *R*_Σ_ with an average error of 37% across the seven PSFs. Note that computing Γ_Σ_ is faster than calculating the average CRB over all of orientation space (2500 sampling points) by a factor of ~1000. While the polarized standard PSF doesn’t show the best variance Γ_Σ_, its FI (in terms of [**Γ**^−1^]_*jj*_) is large for the first four second moments (Fig. 1(b)), implying its superior sensitivity to these rotational motions. Therefore, the polarized PSF can estimate the in-plane (*xy*) orientations of fluorescent molecules with excellent precision (Fig. S2).

**FIG. 1.**
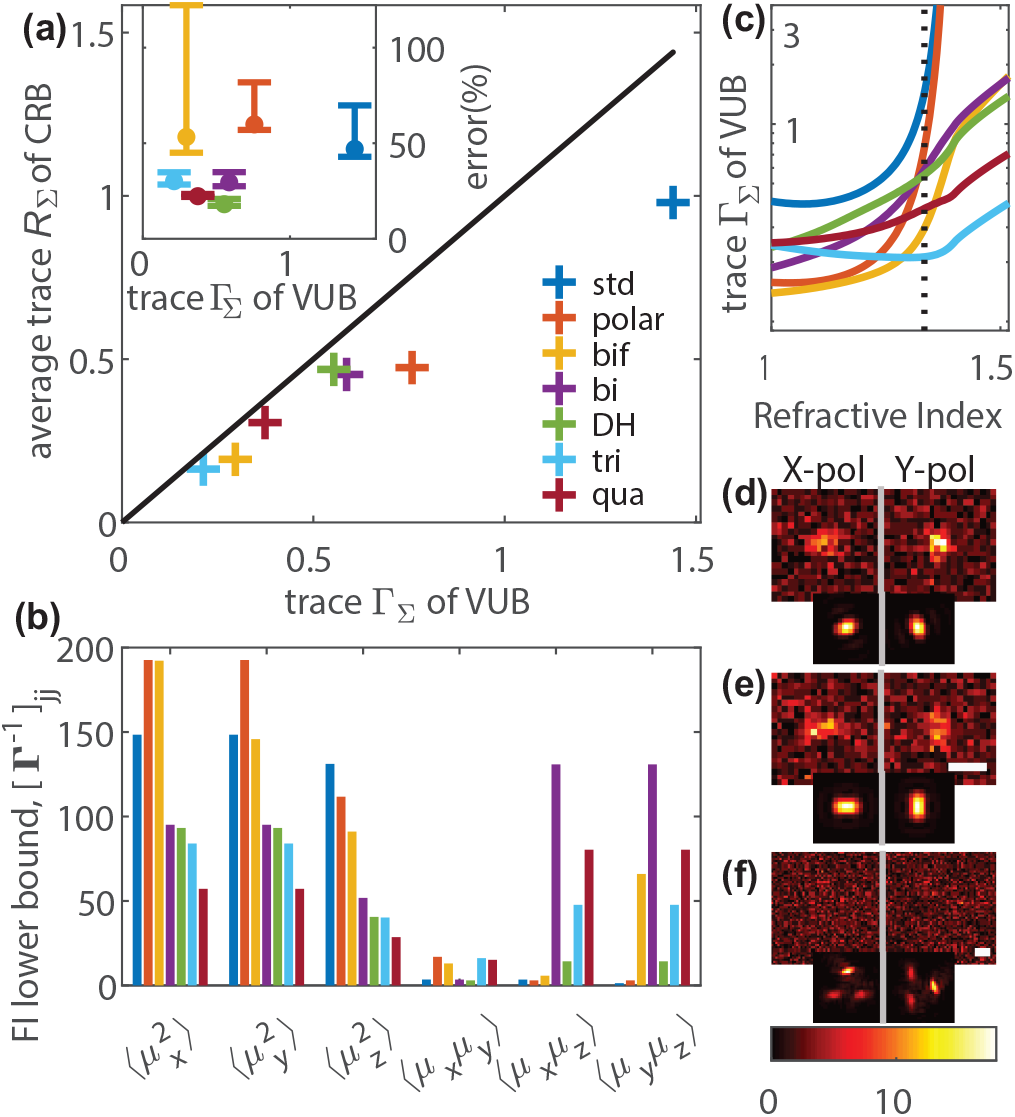
Orientation measurement precision using various point spread functions (PSFs) for 380 signal photons and 2 background photons per pixel. (a) Average (over orientation space) trace *R*_Σ_ of the CRB ***R***, versus the trace Γ_Σ_ of our variance upper bound (VUB) for various PSFs. Blue: unpolarized standard (std), red: polarized standard (polar), yellow: bifocal microscope (bif), purple: bisected (bi), green: double helix (DH), aqua: tri-spot (tri), and maroon: quadrated (qua) PSF. Dark line: Γ_Σ_ = mean of *R*_Σ_. Inset: error of Γ_Σ_ relative to average *R*_Σ_. Bars indicate minimum/maximum error relative to the mean (circle) over all of orientation space. (b) Fisher information (FI) lower bound [**Γ**^*−*1^]_*jj*_ of each orientational second moment *m_j_*. (c) Trace Γ_Σ_ of VUB for various refractive indices. Dotted line: refractive index of water. Simulated orthogonally polarized images of (d) a fixed molecule with orientation (*θ* = 90°, *φ* = 45°) and an isotropic emitter using (e) the polarized PSF and (f) the tri-spot PSF. Insets: normalized noiseless PSF images. Scale bars: 400 nm.

To explore the effect of refractive index (RI) mismatch, we evaluated Γ_Σ_ for sample media ranging from air (RI=1) to oil (RI=1.518). When the RI is low, the polarized standard PSF and bifocal microscope surprisingly show superior measurement performance for the 3D second moments (Fig. 1(c)); these observations also hold for angular orientation parameters (*φ, θ,* Ω) (Fig. S3). This insight is confirmed by our observations that a molecule at an air-glass interface produces basis images ***B**_j_* with vastly superior contrast (energy) compared to one within matched media (Fig. S4). On the other hand, for samples with RI near that of oil, PSF engineering clearly provides increased orientation precision over other approaches. We also quantified the error between VUB and the maximum trace of CRB for a range of RIs (Fig. S5). We observe that the errors are of similar magnitude and are bounded below 60% for all RIs and techniques. Therefore, if VUB predicts that one method is superior to another in terms of measurement variance, then CRB confirms this finding.

In addition to high orientation measurement precision, PSFs must be easily detectable under low SBRs in order to be practical for SMOLM. Using the polarized standard PSF, a fixed molecule (*θ* = 90°, *φ* = 45°) and an isotropic emitter can be distinguished from one another with only 380 photons detected (Figs. 1(d,e), S6). Note that distinguishing these two types of emitters is nearly impossible under matched RI conditions; it is the RI mismatch and the associated supercritical fluorescence collected from fluorophores near the interface (Fig. S7) that enables this remarkable sensitivity. However, the tri-spot PSF, despite its superior VUB, cannot detect the same isotropic emitter due to the splitting of photons into three spots in each polarization channel (Fig. 1(f)). These observations suggest that the polarized standard PSF should be practical for measuring the positions and orientations of single fluorescent molecules precisely and sensitively.

## III. RESOLVING STRUCTURAL HETEROGENEITIES BETWEEN AMYLOID FIBERS

Amyloid aggregates are signatures of various neurodegenerative disorders such as Alzheimer’s and Parkinson’s diseases, and small soluble oligomeric intermediates formed during amyloid fibrillation are typically associated with cytotoxicity [31, 32]. Although many amyloid aggregates share a common cross-*β* sheet structural motif that is an assembly of identical copies of amyloid peptides, oligomers and fibrils are highly heterogeneous in terms of their structure and size [33, 34]. Interestingly, not all aggregation intermediates are equally toxic [35]. Here, we explore using SMOLM with the polarized standard PSF and Nile red (NR) for detecting the structural heterogeneity of amyloid fibrils. NR has been used as a surface hydrophobicity sensor for individual protein aggregates by measuring its emission wavelength while transiently and spontaneously bound to fibrils and proto-fibrils [36, 37]. In this paper, we characterize NR’s binding orientation as a method of mapping the organization of *β*-sheets within amyloid aggregates.

We prepared fibrils of amyloid-*β* peptide (A*β*42) adhered to cleaned glass-bottom imaging chambers as described previously [12] (Supplementary Material 2A). By adding 50 nM NR and illuminating with a circularly polarized 561-nm laser, we captured isolated fluorescence bursts corresponding to individual NR molecules transiently bound to the amyloid aggregates [12, 38] (Movie S1). Note that the circularly polarized illumination preferentially pumps NR molecules lying mostly parallel to the coverslip–precisely those molecules for which the polarized PSF has the best sensitivity for measuring orientations and wobbling areas (Fig. S2). The brightness (*s*), 2D position (*x, y*) and second-moment vector (***m***) of individual NRs within each frame were jointly estimated by using a sparsity-promoting maximum-likelihood estimator [13, 14], followed by a weighted least-square estimator for projecting the estimated second moments to angular coordinates (*φ, θ,* Ω) (Supplementary Material 2D). For simulated images of NR using the polarized PSF, the estimator shows good accuracy and precision for both localization and orientation measurements, especially for molecules approximately parallel to the coverslip (Fig. S8-S13).

The concentrated photon distribution of the polarized PSF enables our estimator to detect relatively dim molecules, providing > 1.4 localizations/nm on amyloid aggregates over ~3 min measurement time (Table S1, Dataset 1 [30]). Fig. 2(a) shows a super-resolved fibril network. We estimate our localization precision to be 9 nm (std. dev., Fig. S9, Table S1). NR molecules exhibit strikingly different polarized PSFs depending on the long axis of the underlying fibrils (Fig. S15). Our orientation measurements show mostly in-plane polar angles (*θ* ≈ 90°, Fig. S16) and azimuthal orientations (*φ*) aligned to the long axis of amyloid fibrils (Figs. 2(b)-2(f), Dataset 1 [30]). These observations confirm that NR molecules preferentially bind to surfaces of *β*-sheets with their transition dipoles parallel to the fibril backbones (Fig. 2(b) inset), similar to the binding behavior of Thioflavin T [39, 40]. NR may bind transiently to grooves formed by adjacent amino acid side chains [41].

**FIG. 2.**
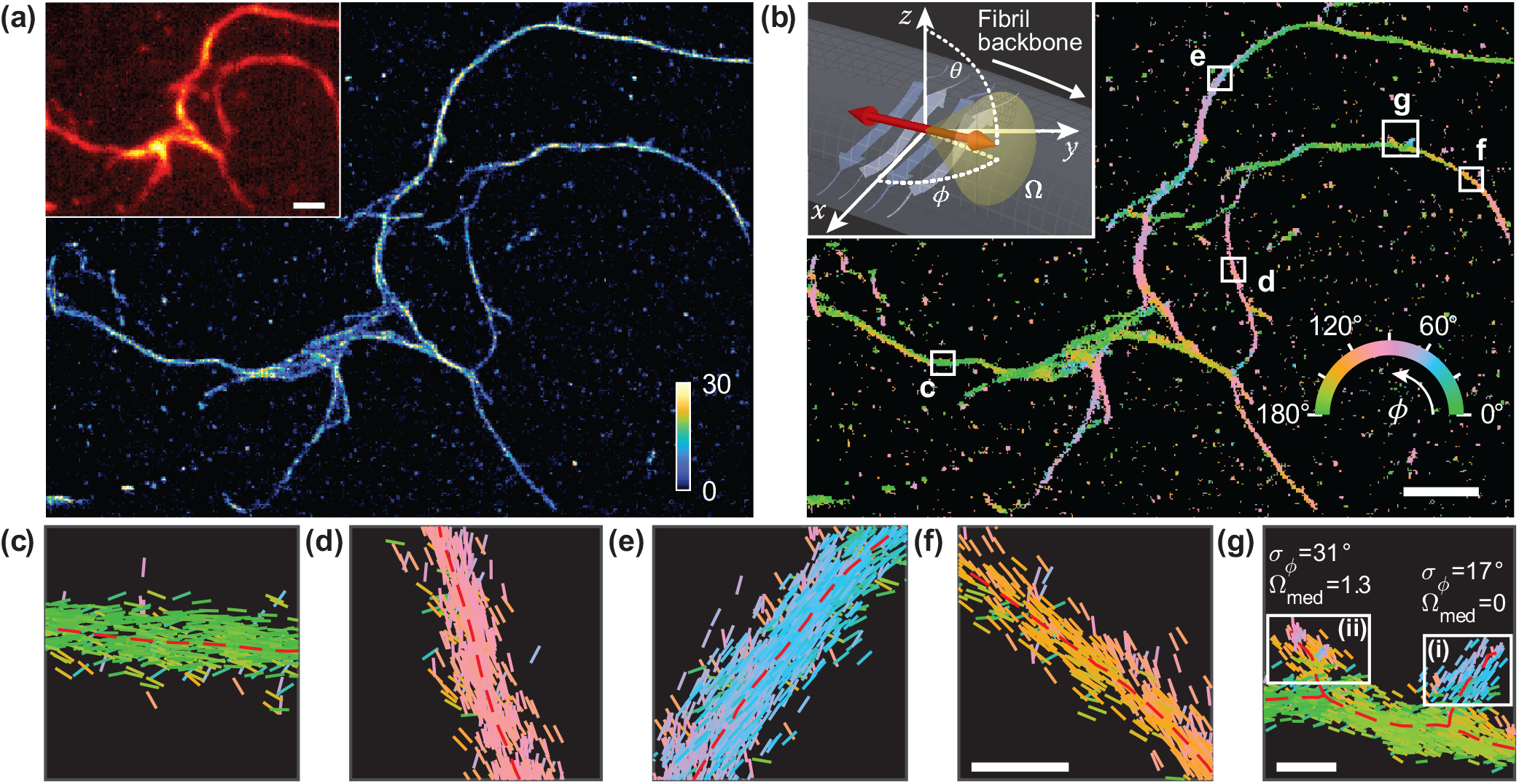
Transient amyloid binding (TAB) SMLM and SMOLM images acquired using Nile red (NR). (a) SMLM image of a network of A*β*42 fibrils. Color bar: localizations per bin (20 × 20 nm^2^). Inset: diffraction-limited image. (b) TAB SMOLM image, color-coded according to the mean azimuthal (*φ*) orientation of NR molecules measured within each bin. Inset: Main binding mode of NR to *β*-sheets, i.e., dipole moments aligned mostly parallel to the long axis of a fibril (its backbone). (c-g) All individual orientation measurements localized along fibril backbones within the white boxes in (b). The lines are oriented and color-coded according to the direction of the estimated *φ* angle. Red dashed lines depict fibril backbones estimated from the SMLM image. Scale bars: (a,b) 1 μm, (f,g) 100 nm. Orientation-localization data are available in Dataset 1 [30].

Although we observed most NR orientations to be parallel to the fibril backbones, these orientations are not always aligned with one another, even in neighboring structures. For example, two small aggregation limbs shown in Fig. 2(g) exhibit substantial differences in the standard deviation of azimuthal orientations (*σ_φ_*) and median wobbling area (Ω_med_), despite looking quite similar in the standard SMLM image. We speculate that this variation arises from structural dissimilarity between these regions, i.e., one of the limbs (Fig. 2(g)(i)) could be a relatively new deposit with disordered binding sites (leading to diverse mean orientations and large wobble), while the other (Fig. 2(g)(ii)) is a relatively well-ordered secondary nucleation of a new fibril branch [34, 42]. These structural details of amyloid aggregates cannot be detected by standard SMLM.

To further correlate the orientations of single-molecule probes with the structural organization of amyloid aggregates, we imaged an amyloid aggregate containing obvious fibril bundles (Fig. 3(a), Dataset 1 [30]). The intertwined structures are evidenced by their branching architecture and their at least twice larger width (purple) compared to thinner structures (green) in the field of view (Fig. S19). We observed NR to be mostly parallel to the backbone (Figs. 3(b)(i), 3(c)(i), and Movie S2) of the thin fibrils, whereas thicker bundles cause NR to bind with relatively disordered *φ* angles (Figs. 3(b)(ii), 3(c)(ii), and Movie S2). Since NR binds to specific grooves along the stacked *β*-sheet aggregate, these orientation measurements indicate highly aligned *β*-sheets within the thin fibrils. We consistently observed similarly disordered *φ* orientations of NR on thicker aggregates compared to thinner structures (Figs. S21 and S22). These disordered *φ* angles show that the *β*-sheets within bundles of amyloid fibrils exhibit a complex and heterogeneous architecture. We note that the wobbling area Ω, which can indicate the binding strength between NR and the surrounding *β*-sheet grooves, also exhibits obvious differences between the two regions. We observed small Ω values on the thin fibril (Figs. 3(d)(i), S23(a), and S23(b)) and large wobbling on the fibril bundle (Figs. 3(d)(ii), S23(a), and S23(c)) that represent tight and loose binding of NR molecules to each local region, respectively. From these measurements, we speculate that fibril bundling not only produces relatively disordered *β*-sheet orientations due to the entwining of multiple fibers but also increases dramatically the local density of binding sites for NR, thereby causing large NR wobbling during a single fluorescence burst. Note that we did not observe significantly different burst times for NR on thin fibers (*τ*_on_ = 13 ms in Fig. 3(b)(i)) versus disordered fibrillar bundles (*τ*_on_ = 11 ms in Fig. 3(b)(ii)).

**FIG. 3.**
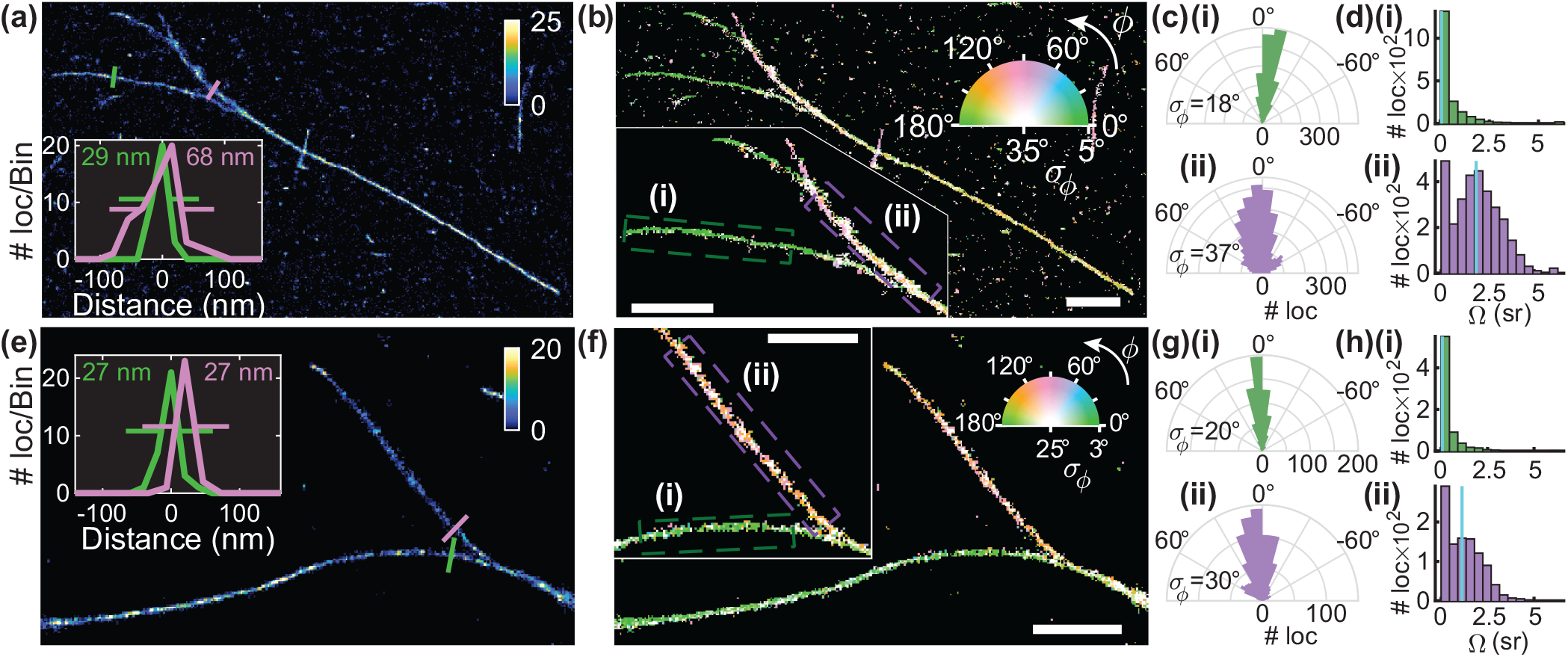
Structural heterogeneity of A*β*42 fibrils revealed by TAB SMOLM imaging. (a) SMLM image of fibril bundles. Color bar: localizations per bin (20 × 20 nm^2^). Inset: fibril cross-sections at the locations denoted by green and purple lines with measured full-width at half-maximum (FWHM) thicknesses. (b) TAB SMOLM image corresponding to (a), color-coded according to the mean (*φ*) and standard deviation (*σ_φ_*) of the azimuthal orientation measured within each bin. Inset: zoomed (i) thin and (ii) thick fibril regions isolated from background structures. (c) Histograms of measured azimuthal orientations relative to the fibril backbone within the regions denoted in (b), showing standard deviations (*σ_φ_*) of (i) 18° and (ii) 37°. (d) Measured wobbling areas (Ω) corresponding to the localizations in (c), yielding median wobbling areas (Ω_med_, cyan) of (i) 0.07 sr and (ii) 1.89 sr. (e-h) TAB SMLM and SMOLM images of another fibril field of view, as in (a-d). Although fibril regions (i) and (ii) in the inset of (f) show little difference in apparent width, the measured orientation distributions contain significant differences. The standard deviations of azimuthal angles *σ_φ_* are (i) 20° vs (ii) 30°, and the median wobbling areas (Ω_med_, cyan) are (i) 0.07 sr vs (ii) 1.12 sr. Scale bars: 1 μm. Orientation-localization data are available in Dataset 1 [30].

Interestingly, the local structural heterogeneity in amyloid fibrils detected by SMOLM is not always associated with thicker fibrils in standard SMLM. We observed similar disordered *φ* values and large Ω estimates from a fibril in another field of view (Fig 3(e)). Although the two fibril strands have similar thickness, the estimated *φ* and Ω distributions show noticeable differences (Fig. 3(f)-3(h)). These measurements suggest the fibril in Fig. 3(f)(ii) may consist of more disorganized assemblies of *β*-sheets compared to those of Fig. 3(f)(i). Similar to the small aggregates shown in Fig. 2(g), the local heterogeneity cannot be resolved by conventional SMLM.

## IV. CONCLUSION

Here, we propose and characterize a new variance upper bound (VUB) for efficiently quantifying the sensitivity of any PSF for measuring the orientation and wobble of dipole-like emitters. Since the orientational second moment vector ***m*** is six dimensional, VUB greatly accelerates the comparison of various PSFs for any specific imaging scenario. While VUB overestimates the mean variance of an orientation measurement by 37%, computing the VUB is ~1000 times faster than evaluating the average CRB across orientation space with sufficient sampling. We stress that this computational speedup of measuring global performance makes VUB a useful design and optimization tool, but CRB is still useful for quantitatively characterizing measurement performance for specific molecular orientations ***m***. Using VUB, we discover that imaging dipoles near a refractive index interface enables the simple polarized standard PSF to robustly measure orientational dynamics, particularly within the xy plane, that cannot be detected in matched media.

Using the polarized standard PSF, our SMOLM images of amyloid aggregates confirm that the main binding orientation of NR is mostly parallel to the long axis of the fibrils. In cases where multiple fibrils are intertwined, TAB SMOLM detects heterogeneities in the stacking of *β*-sheets via larger variations in NR orientations and increased wobbling angles. In particular, SMOLM detects structural disorder along the fibrillar network which cannot be resolved by simple SMLM analysis of fibrillar width or NR binding time.

Simultaneous imaging of single-molecule positions and orientations in SMOLM allows scientists to quantify local structural features on a nanometer scale without ensemble averaging. In particular, measuring the full spatial configuration (i.e., positions and orientations) of molecules provides unprecedented insight into the density and orientations of binding sites for NR on aggregate surfaces that aren’t revealed with location information alone. These distributions could be a signature of how the stacked *β*-sheets are organized within fibrils and other aggregate morphologies. Since our analysis shows that no single PSF measures all combinations of orientational second moments with optimal precision, the superior speed and sufficient accuracy of VUB may be useful for adaptively optimizing PSFs for a particular sample or field of view during a SMOLM experiment. Such optimization would need to be performed in near real-time to obtain the best-possible measurement precision while minimizing photobleaching and photodamage within the specific sample of interest.

Currently, no methodology can visualize the nanoscale structural configuration of amyloid aggregates as they organize over time at the single particle level [34]. In the future, SMOLM combined with transient binding of amyloid-sensitive fluorogenic molecules could become a powerful tool for understanding how structure affects various aggregation processes, such as “stop-and-go” and polarized aggregation [42–44] as well as the switching of aggregation modes [45], with nanoscale resolution over periods of hours to days [12, 38]. Time-lapse SMOLM imaging of amyloidophilic dye-binding sites on aggregate surfaces could provide insight into amyloid assembly mechanisms [44, 46] by connecting the energy landscapes of these pathways to *β*-sheet organization over time. Finally, TAB SMOLM imaging of amyloid aggregates on model membranes and neurons may provide links between aggregate structure and toxicity necessary for developing effective therapeutics against amyloidosis.

## Funding Information

National Science Foundation (NSF) (ECCS-1653777); National Institute of General Medical Sciences of the National Institutes of Health (R35GM124858).

## Acknowledgments

The authors thank Jan Bieschke, Jin Lu, Yuanzi Sun and Kevin Spehar for helpful suggestions and comments.

## Disclosures

The authors declare no conflicts of interest.

See Supplement 1 for supporting content. Dataset 1 is available via OSF [30] and by request.

## Notes

https://osf.io/pe3qu/?view_only=081206495472426889c1055f21971e9a

### Competing Interest Statement

The authors have declared no competing interest.

### Summary of Updates

Updated VUB analysis: first- and second-order orientational moments, comparison to CRB, additional methods and sample configurations; additional citations to related literature

## References

[1] H. Sosa, E. J. Peterman, W. E. Moerner, and L. S. Goldstein, ADP-induced rocking of the kinesin motor domain revealed by single-molecule fluorescence polarization microscopy, Nature Structural Biology 8, 540 (2001).

[2] E. J. Peterman, H. Sosa, L. S. Goldstein, and W. Moerner, Polarized Fluorescence Microscopy of Individual and Many Kinesin Motors Bound to Axonemal Microtubules, Biophysical Journal 81, 2851 (2001).

[3] L. G. Lippert, T. Dadosh, J. A. Hadden, V. Karnawat, B. T. Diroll, C. B. Murray, E. L. F. Holzbaur, K. Schulten, S. L. Reck-Peterson, and Y. E. Goldman, Angular measurements of the dynein ring reveal a stepping mechanism dependent on a flexible stalk, Proceedings of the National Academy of Sciences 114, E4564 (2017).

[4] T. Ha, J. Glass, T. Enderle, D. S. Chemla, and S. Weiss, Hindered Rotational Diffusion and Rotational Jumps of Single Molecules, Physical Review Letters 80, 2093 (1998).

[5] A. S. Backer, A. S. Biebricher, G. A. King, G. J. L. Wuite, I. Heller, and E. J. G. Peterman, Single-molecule polarization microscopy of DNA intercalators sheds light on the structure of S-DNA, Science Advances 5, eaav1083 (2019).

[6] B. Dong, Y. Pei, N. Mansour, X. Lu, K. Yang, W. Huang, and N. Fang, Deciphering nanoconfinement effects on molecular orientation and reaction intermediate by single molecule imaging, Nature Communications 10, 4815 (2019).

[7] M. P. Backlund, M. D. Lew, A. S. Backer, S. J. Sahl, and W. Moerner, The role of molecular dipole orientation in single-molecule fluorescence microscopy and implications for super-resolution imaging, ChemPhysChem 15, 587 (2014).

[8] R. J. Ober, S. Ram, and E. S. Ward, Localization accuracy in single-molecule microscopy., Biophysical journal 86, 1185 (2004).

[9] G. Grover, K. DeLuca, S. Quirin, J. DeLuca, and R. Piestun, Super-resolution photon-efficient imaging by nanometric double-helix point spread function localization of emitters (SPINDLE), Optics Express 20, 26681 (2012).

[10] Y. Shechtman, S. J. Sahl, A. S. Backer, and W. E. Moerner, Optimal point spread function design for 3D imaging, Physical Review Letters 113, 1 (2014).

[11] D. Patra, I. Gregor, and J. Enderlein, Image analysis of defocused single molecule images for three dimensional molecular orientation studies, J. Phys. Chem. A 108, 6836 (2004).

[12] K. Spehar, T. Ding, Y. Sun, N. Kedia, J. Lu, G. R. Nahass, M. D. Lew, and J. Bieschke, Super-resolution Imaging of Amyloid Structures over Extended Times by Using Transient Binding of Single Thioflavin T Molecules, ChemBioChem 19, 1944 (2018).

[13] H. Mazidi, E. S. King, O. Zhang, A. Nehorai, and M. D. Lew, Dense Super-Resolution Imaging of Molecular Orientation Via Joint Sparse Basis Deconvolution and Spatial Pooling, in IEEE 16th International Symposium on Biomedical Imaging (ISBI), 325–329 (2019).

[14] H. Mazidi, T. Ding, and M. D. Lew, https://github.com/Lew-Lab/RoSE-O.

[15] D. L. Sackett and J. Wolff, Nile red as a polarity-sensitive fluorescent probe of hydrophobic protein surfaces, Analytical Biochemistry 167, 228 (1987).

[16] A. Sharonov and R. M. Hochstrasser, Wide-field subdiffraction imaging by accumulated binding of diffusing probes, Proceedings of the National Academy of Sciences 103, 18911 (2006).

[17] L. Novotny and B. Hecht, Principles of Nano-Optics (Cambridge University Press, 2006).

[18] T. Chandler, H. Shroff, R. Oldenbourg, and P. La Rivière, Spatio-angular fluorescence microscopy i. basic theory, JOSA A 36, 1334 (2019).

[19] T. Chandler, H. Shroff, R. Oldenbourg, and P. La Rivière, Spatio-angular fluorescence microscopy ii. paraxial 4f imaging, JOSA A 36, 1346 (2019).

[20] S. Stallinga, Effect of rotational diffusion in an orientational potential well on the point spread function of electric dipole emitters, JOSA A 32, 213 (2015).

[21] A. S. Backer and W. Moerner, Extending single-molecule microscopy using optical fourier processing, The Journal of Physical Chemistry B 118, 8313 (2014).

[22] A. S. Backer and W. Moerner, Determining the rotational mobility of a single molecule from a single image: a numerical study, Optics express 23, 4255 (2015).

[23] M. Böhmer and J. Enderlein, Orientation imaging of single molecules by wide-field epifluorescence microscopy, JOSA B 20, 554 (2003).

[24] M. A. Lieb, J. M. Zavislan, and L. Novotny, Single-molecule orientations determined by direct emission pattern imaging, JOSA B 21, 1210 (2004).

[25] A. Agrawal, S. Quirin, G. Grover, and R. Piestun, Limits of 3d dipole localization and orientation estimation for single-molecule imaging: towards green’s tensor engineering, Optics express 20, 26667 (2012).

[26] A. S. Backer, M. P. Backlund, A. R. von Diezmann, S. J. Sahl, and W. Moerner, A bisected pupil for studying single-molecule orientational dynamics and its application to three-dimensional super-resolution microscopy, Applied physics letters 104, 193701 (2014).

[27] M. P. Backlund, M. D. Lew, A. S. Backer, S. J. Sahl, G. Grover, A. Agrawal, R. Piestun, and W. Moerner, Simultaneous, accurate measurement of the 3d position and orientation of single molecules, Proceedings of the National Academy of Sciences 109, 19087 (2012).

[28] O. Zhang, J. Lu, T. Ding, and M. D. Lew, Imaging the three-dimensional orientation and rotational mobility of fluorescent emitters using the tri-spot point spread function, Applied physics letters 113, 031103 (2018).

[29] A. S. Backer, M. P. Backlund, M. D. Lew, and W. E. Moerner, Single-molecule orientation measurements with a quadrated pupil, Optics letters 38, 1521 (2013).

[30] M. D. Lew, T. Ding, and T. Wu, https://osf.io/pe3qu/?view_only=081206495472426889c1055f21971e9a.

[31] E. Cohen, J. Bieschke, R. M. Perciavalle, J. W. Kelly, and A. Dillin, Opposing Activities Protect Against Age-Onset Proteotoxicity, Science 313, 1604 (2006).

[32] M. Serra-Batiste, M. Ninot-Pedrosa, M. Bayoumi, M. Gairí, G. Maglia, and N. Carulla, A*β*42 assembles into specific *β*-barrel pore-forming oligomers in membrane-mimicking environments, Proceedings of the National Academy of Sciences 113, 10866 (2016).

[33] L. M. Young, L.-H. Tu, D. P. Raleigh, A. E. Ashcroft, and S. E. Radford, Understanding co-polymerization in amyloid formation by direct observation of mixed oligomers, Chemical Science 8, 5030 (2017).

[34] M. G. Iadanza, M. P. Jackson, E. W. Hewitt, N. A. Ranson, and S. E. Radford, A new era for understanding amyloid structures and disease, Nature Reviews Molecular Cell Biology 19, 755 (2018).

[35] G. Fusco, S. W. Chen, P. T. F. Williamson, R. Cascella, M. Perni, J. A. Jarvis, C. Cecchi, M. Vendruscolo, F. Chiti, N. Cremades, L. Ying, C. M. Dobson, and A. De Simone, Structural basis of membrane disruption and cellular toxicity by *α*-synuclein oligomers, Science 358, 1440 (2017).

[36] M. N. Bongiovanni, J. Godet, M. H. Horrocks, L. Tosatto, A. R. Carr, D. C. Wirthensohn, R. T. Ranasinghe, J. E. Lee, A. Ponjavic, J. V. Fritz, C. M. Dobson, D. Klenerman, and S. F. Lee, Multi-dimensional super-resolution imaging enables surface hydrophobicity mapping, Nature Communications 7, 10.1038/ncomms13544 (2016).

[37] J.-E. Lee, J. C. Sang, M. Rodrigues, A. R. Carr, M. H. Horrocks, S. De, M. N. Bongiovanni, P. Flagmeier, C. M. Dobson, D. J. Wales, S. F. Lee, and D. Klenerman, Mapping Surface Hydrophobicity of *α*-Synuclein Oligomers at the Nanoscale, Nano Letters 18, 7494 (2018).

[38] T. Ding, K. Spehar, J. Bieschke, and M. D. Lew, Long-term, super-resolution imaging of amyloid structures using transient amyloid binding microscopy, in Single Molecule Spectroscopy and Superresolution Imaging XII, 108840J, edited by I. Gregor, Z. K. Gryczynski, and F. Koberling (SPIE, 2019).

[39] H. A. Shaban, C. A. Valades-Cruz, J. Savatier, and S. Brasselet, Polarized super-resolution structural imaging inside amyloid fibrils using Thioflavine T, Scientific Reports 7, 12482 (2017).

[40] J. A. Varela, M. Rodrigues, S. De, P. Flagmeier, S. Gandhi, C. M. Dobson, D. Klenerman, and S. F. Lee, Optical Structural Analysis of Individual *α*-Synuclein Oligomers, Angewandte Chemie International Edition 57, 4886 (2018).

[41] M. Biancalana and S. Koide, Molecular mechanism of Thioflavin-T binding to amyloid fibrils, Biochimica et Biophysica Acta - Proteins and Proteomics 1804, 1405 (2010).

[42] C. B. Andersen, H. Yagi, M. Manno, V. Martorana, T. Ban, G. Christiansen, D. E. Otzen, Y. Goto, and C. Rischel, Branching in Amyloid Fibril Growth, Biophysical Journal 96, 1529 (2009).

[43] L. J. Young, G. S. Kaminski Schierle, and C. Kaminski, Imaging A*β*(1–42) fibril elongation reveals strongly polarised growth and growth incompetent states, Phys. Chem. Chem. Phys. 19, 27987 (2017).

[44] A. K. Buell, The growth of amyloid fibrils: rates and mechanisms, Biochemical Journal 476, 2677 (2019).

[45] T. Watanabe-Nakayama, K. Ono, M. Itami, R. Takahashi, D. B. Teplow, and M. Yamada, High-speed atomic force microscopy reveals structural dynamics of amyloid *β* 1–42 aggregates, Proceedings of the National Academy of Sciences 113, 5835 (2016).

[46] T. Eichner and S. E. Radford, A Diversity of Assembly Mechanisms of a Generic Amyloid Fold, Molecular Cell 43, 8 (2011).

