## Supplement 1 for "Single-molecule orientation localization microscopy for resolving structural heterogeneities between amyloid fibrils"

### Abstract

This document provides supplementary information to "Single-molecule orientation localization microscopy for resolving structural heterogeneities between amyloid fibrils," offering details on the derivation of VUB, VUB evaluated for first- and second-order orientational moments, TAB experimental methods, the accuracy and precision of the SMOLM estimator, and additional SMOLM images and measurements.

#### CONTENTS

List of Figures 3

List of Tables 4

I. Bounding Fisher information and measurement variance 5
A. Variance upper bound 5
B. Variance upper bound for subsets of orientational second-moments 6
C. Variance upper bound for first-moment cone angles 7

II. Experimental methods 8
A. Preparation of amyloid aggregates 8
B. Optical instrumentation and imaging procedure 9
C. Background estimation and dual-channel registration 9
D. Localization and orientation estimation of single molecules 10
E. Selection of fibril region of interest and measurement of fibril width 12
F. Synthetic data 12

References 37

* These authors contributed equally to this work

†

S5 Comparison of CRB and VUB for emitters at various refractive index interfaces 18
S6 Resolvability of a fixed molecule versus an isotropic emitter using our estimator 19
S7 Orientation measurement precision for emitters at various depths within water 20
S8 Cramér–Rao bounds for estimating the 3D position of a single molecule using the polarized and unpolarized standard PSFs . . . . . . . . . . . . . . . . . . . . . . . . . . . . 21
S9 SMOLM measurement precision for position and signal . . . . . . . . . . . . . . . . . . 22
S10 SMOLM measurement bias for position and signal . . . . . . . . . . . . . . . . . . . . . . 23
S11 SMOLM measurement precision for orientation . . . . . . . . . . . . . . . . . . . . . . . . . 24
S12 SMOLM measurement bias for orientation . . . . . . . . . . . . . . . . . . . . . . . . . . . . 25
S13 SMOLM orientational measurement bias for isotropic emitters . . . . . . . . . . . . . 26
S14 Orientation and wobbling area measurements of fluorescent beads . . . . . . . . . . 27
S15 Fluorescence images of single Nile red molecules on fibril backbones with various orientations . . . . . . . . . . . . . . . . . . . . . . . . . . . . . . . . . . . . . . . . . . . . . . . . 28
S16 Estimated polar orientations $\theta$ of Nile red molecules on fibrils. . . . . . . . . . . . . . 28
S17 Fibril region of interest and backbone ridge detection . . . . . . . . . . . . . . . . . . . . 29
S18 Photophysics of Nile red transient amyloid binding microscopy. . . . . . . . . . . . . 30
S19 Measurements of fibril widths . . . . . . . . . . . . . . . . . . . . . . . . . . . . . . . . . . . . . . . 31
S20 Correlation between estimates of azimuthal orientation and wobbling area . . . 32
S21 Additional fibril analysis I . . . . . . . . . . . . . . . . . . . . . . . . . . . . . . . . . . . . . . . . . 33
S22 Additional fibril analysis II. . . . . . . . . . . . . . . . . . . . . . . . . . . . . . . . . . . . . . . . . 34
S23 Estimated wobbling area of Nile red molecules on amyloid fibrils . . . . . . . . . . . 35

**LIST OF TABLES**

S1 Localization numbers, localization density, photons detected per localization, background photons per pixel, and burst on-time of Nile Red molecules on amyloid structures . . . . . 36

### I. BOUNDING FISHER INFORMATION AND MEASUREMENT VARIANCE

We define the Fisher information $\boldsymbol{J}$ associated with estimating the six orientational second moments $\boldsymbol{m} = \left[\langle\mu_x^2\rangle, \langle\mu_y^2\rangle, \langle\mu_z^2\rangle, \langle\mu_x\mu_y\rangle, \langle\mu_x\mu_z\rangle, \langle\mu_y\mu_z\rangle\right]^T$ as

$$\boldsymbol{J} = \sum_{i=1}^{n} \frac{1}{I_i} \nabla I_i^T \nabla I_i, \tag{S1}$$

where the subscript $i$ denotes the $i^{\text{th}}$ element of $\boldsymbol{I} \in \mathbb{R}^n$, the expected photon distribution captured by a camera with $n$ pixels. Our forward imaging model allows us to derive a simple expression for $\boldsymbol{J}$ as:

$$\boldsymbol{I} = s\begin{bmatrix}\boldsymbol{B}_{xx} & \boldsymbol{B}_{yy} & \boldsymbol{B}_{zz} & \boldsymbol{B}_{xy} & \boldsymbol{B}_{xz} & \boldsymbol{B}_{yz}\end{bmatrix}\boldsymbol{m} + \boldsymbol{b} \tag{S2}$$

$$\begin{aligned} \implies \nabla \boldsymbol{I}_i &= \begin{bmatrix}\frac{\partial I_i}{\partial m_1} & \frac{\partial I_i}{\partial m_2} & \frac{\partial I_i}{\partial m_3} & \frac{\partial I_i}{\partial m_4} & \frac{\partial I_i}{\partial m_5} & \frac{\partial I_i}{\partial m_6}\end{bmatrix} \\ &= s\boldsymbol{B}_i \end{aligned} \tag{S3}$$

$$\begin{aligned} \implies \boldsymbol{J} &= \sum_{i=1}^{n} \frac{s^2}{I_i} \boldsymbol{B}_i^T \boldsymbol{B}_i \\ &\triangleq \boldsymbol{A}^T\boldsymbol{A}, \end{aligned} \tag{S4}$$

where $\boldsymbol{B}_i$ denotes the $i^{\text{th}}$ row of $\boldsymbol{B} \in \mathbb{R}^{n\times 6}$ and the rows of $\boldsymbol{A}$ are given by $\boldsymbol{A}_i = s\boldsymbol{B}_i/\sqrt{s\boldsymbol{B}_i\boldsymbol{m} + b_i}$.

#### A. Variance upper bound

To derive the variance upper bound (VUB), we first introduce an $n \times n$ matrix $\boldsymbol{C}$ as

$$\boldsymbol{C} = \begin{bmatrix} \frac{\sqrt{s\|\boldsymbol{B}_1\|+b_1}}{\sqrt{s\boldsymbol{B}_1\boldsymbol{m}+b_1}} & 0 & \dots & 0 \\ 0 & \frac{\sqrt{s\|\boldsymbol{B}_2\|+b_2}}{\sqrt{s\boldsymbol{B}_2\boldsymbol{m}+b_2}} & \dots & 0 \\ \vdots & \vdots & \ddots & \vdots \\ 0 & 0 & \dots & \frac{\sqrt{s\|\boldsymbol{B}_n\|+b_n}}{\sqrt{s\boldsymbol{B}_n\boldsymbol{m}+b_n}} \end{bmatrix}, \tag{S5}$$

where $n$ is the number of pixels in the image $\boldsymbol{I}$. The Cauchy-Schwarz inequality implies $\boldsymbol{B}_i\boldsymbol{m} \le \|\boldsymbol{B}_i\|\,\|\boldsymbol{m}\|$. Since $\|\boldsymbol{\mu}\| = 1$ for any SM transition dipole, $\|\boldsymbol{m}\| \le 1$ for all possible dipole orientations. Therefore, we may write an orientation-independent bound using $\boldsymbol{B}_i\boldsymbol{m} \le \|\boldsymbol{B}_i\|$, and $\boldsymbol{C}$ is a diagonal matrix whose entries are greater than or equal to 1. We rewrite $\boldsymbol{J}$ in terms of $\boldsymbol{C}$ as

$$\boldsymbol{J} = \hat{\boldsymbol{A}}^T\boldsymbol{C}^T\boldsymbol{C}\hat{\boldsymbol{A}}, \tag{S6}$$

where the rows of $\hat{\boldsymbol{A}}$ are given by

$$\hat{\boldsymbol{A}}_i = \frac{s\boldsymbol{B}_i}{\sqrt{s\left\|\boldsymbol{B}_i\right\| + b_i}}. \tag{S7}$$

Therefore, a lower bound on FI for measuring a specific orientational second moment $m_j$ can be represented as

$$J_{jj} \geq \left[\left(\hat{\boldsymbol{A}}^T\hat{\boldsymbol{A}}\right)\right]_{jj} = \left[\boldsymbol{\Gamma}^{-1}\right]_{jj}, \tag{S8}$$

where the subscript $jj$ represents the $j^{\text{th}}$ diagonal element of the matrix. We term $\boldsymbol{\Gamma}$ the VUB matrix and define it as

$$\boldsymbol{\Gamma} \triangleq \left(\hat{\boldsymbol{A}}^T\hat{\boldsymbol{A}}\right)^{-1}. \tag{S9}$$

We next prove that VUB is a global upper bound of the Cramér–Rao bound (CRB) for all possible orientational second moments. The CRB matrix $\boldsymbol{R}$ can be represented as

$$\begin{aligned}\boldsymbol{R} = \boldsymbol{J}^{-1} &= \left(\boldsymbol{\Gamma}^{-\mathbf{1}} + \left(\boldsymbol{J} - \boldsymbol{\Gamma}^{-\mathbf{1}}\right)\right)^{-1} \\ &= \boldsymbol{\Gamma} - \boldsymbol{\Gamma}\left(\boldsymbol{J} - \boldsymbol{\Gamma}^{-\mathbf{1}}\right)\boldsymbol{J}^{-1},\end{aligned} \tag{S10}$$

where both $\boldsymbol{\Gamma}$ and $\boldsymbol{J}^{-1}$ are positive semi-definite. We may write

$$\boldsymbol{J} - \boldsymbol{\Gamma}^{-\mathbf{1}} = \left(\hat{\boldsymbol{A}}^T\boldsymbol{\Lambda}\right)\left(\hat{\boldsymbol{A}}^T\boldsymbol{\Lambda}\right)^T, \tag{S11}$$

where $\boldsymbol{\Lambda} = \left(\boldsymbol{C}^T\boldsymbol{C} - \boldsymbol{E}\right)^{1/2}$ and $\boldsymbol{E}$ is the identity matrix; therefore, $(\boldsymbol{J} - \boldsymbol{\Gamma}^{-\mathbf{1}})$ is also positive semi-definite. Recall that the eigenvalues of the product of two positive-definite matrices are non-negative [1]. Examining $\boldsymbol{\Gamma}\left(\boldsymbol{J} - \boldsymbol{\Gamma}^{-\mathbf{1}}\right)\boldsymbol{J}^{-1}$, which is the product of three positive semi-definite matrices, we find that both its eigenvalues and diagonal entries are non-negative.

We therefore have

$$R_{jj} \leq \Gamma_{jj}, \tag{S12}$$

that is, each diagonal element of the VUB matrix $\boldsymbol{\Gamma}$ is an upper bound on the corresponding element of the CRB matrix $\boldsymbol{R}$. We may calculate the largest possible measurement variance of any efficient unbiased estimator for any possible molecular orientation trajectory (i.e., second moment vector $\boldsymbol{m}$) using Eqn. (S9).

#### B. Variance upper bound for subsets of orientational second-moments

Certain samples, such as lipid membranes or the amyloid fibers in this work, create special geometries that limit or constrain the range of single-molecule (SM) orientations to be measured. In this case, we can use the limited photon budget more effectively by focusing on measuring only a subset of second moments $\boldsymbol{m}$.

We next modify our VUB matrix for in-plane oriented molecules (i.e., perpendicular to the optical axis or $\langle \mu_z \rangle = 0$), where $\langle \cdot \rangle$ represents temporal averaging over a camera frame. For in-plane oriented molecules, only 2D second moments $\boldsymbol{m}_{\text{2D}} = [\langle \mu_x^2 \rangle, \langle \mu_y^2 \rangle, \langle \mu_z^2 \rangle, \langle \mu_x \mu_y \rangle]$ and the 2D basis matrix $\boldsymbol{B}_{\text{2D}} = [\boldsymbol{B}_{xx}, \boldsymbol{B}_{yy}, \boldsymbol{B}_{zz}, \boldsymbol{B}_{xy}]$ are of interest for characterizing estimation precision. Therefore, we have a 2D CRB matrix $\boldsymbol{R}_{\text{2D}}$ given by

$$\boldsymbol{R}_{\text{2D}} = \boldsymbol{J}_{\text{2D}}^{-1} = \left( \boldsymbol{A}_{\text{2D}}^T \boldsymbol{A}_{\text{2D}} \right)^{-1}, \tag{S13}$$

where the rows of $\boldsymbol{A}_{\text{2D}}$ are given by $\boldsymbol{A}_{\text{2D},i} = s\boldsymbol{B}_{\text{2D},i} / \sqrt{s\boldsymbol{B}_{\text{2D},i}\boldsymbol{m}_{\text{2D}} + b_i}$, and $\boldsymbol{B}_{\text{2D},i}$ is the $i^{th}$ row of $\boldsymbol{B}_{\text{2D}}$.

Similar to the aforementioned (3D) VUB, we define our 2D VUB matrix $\boldsymbol{\Gamma}_{\text{2D}}$ as

$$\boldsymbol{\Gamma}_{\text{2D}} = \left( \hat{\boldsymbol{A}}_{\text{2D}}^T \hat{\boldsymbol{A}}_{\text{2D}} \right)^{-1}, \tag{S14}$$

where

$$\hat{\boldsymbol{A}}_{\text{2D},i} = \frac{s\boldsymbol{B}_{\text{2D},i}}{\sqrt{s \left\| \boldsymbol{B}_{\text{2D},i} \right\| + b_i}}. \tag{S15}$$

#### C. Variance upper bound for first-moment cone angles

To measure molecular orientation using a "wobble-in-a-cone" model, we first estimate the orientational second moments from observed images and then project these values into first-moment space in terms of angles of a cone ($(\phi, \theta, \Omega)$, see Fig. S1, Sec. 2II D). Therefore, the estimation precision of second moments directly determines the estimation precision in angular space.

Here, we derive an angular VUB $\boldsymbol{\Gamma}_{\text{ang}}$ for bounding the angular variance $\left( \sigma_\phi^2, \sigma_\theta^2, \sigma_\Omega^2 \right)$ at each orientation state. We first define a transform matrix $\boldsymbol{Q}$ as

$$\boldsymbol{Q} = \begin{bmatrix} \frac{\partial m_1}{\partial \phi} & \frac{\partial m_2}{\partial \phi} & \frac{\partial m_3}{\partial \phi} & \frac{\partial m_4}{\partial \phi} & \frac{\partial m_5}{\partial \phi} & \frac{\partial m_6}{\partial \phi} \\ \frac{\partial m_1}{\partial \theta} & \frac{\partial m_2}{\partial \theta} & \frac{\partial m_3}{\partial \theta} & \frac{\partial m_4}{\partial \theta} & \frac{\partial m_5}{\partial \theta} & \frac{\partial m_6}{\partial \theta} \\ \frac{\partial m_1}{\partial \Omega} & \frac{\partial m_2}{\partial \Omega} & \frac{\partial m_3}{\partial \Omega} & \frac{\partial m_4}{\partial \Omega} & \frac{\partial m_5}{\partial \Omega} & \frac{\partial m_6}{\partial \Omega} \end{bmatrix}, \tag{S16}$$

that enables the angular VUB $\boldsymbol{\Gamma}_{\text{ang}}$ to be calculated in terms the VUB $\boldsymbol{\Gamma}$ of second-order moments using

$$\boldsymbol{\Gamma}_{\text{ang}} = \left( \boldsymbol{Q} \boldsymbol{\Gamma}^{-1} \boldsymbol{Q}^T \right)^{-1}. \tag{S17}$$

The $j^{\text{th}}$ diagonal element $\Gamma_{\text{ang},jj}$ is an upper bound on the variance of measuring the $j^{\text{th}}$ corresponding angular parameter $[\phi, \theta, \Omega]$.

As a summary measure of performance, we calculate the root-mean-square angular error (RMSAE) $\sigma_k$ to combine variances of measuring $\phi$ and $\theta$ [2]. The RMSAE upper bound $\sqrt{\Gamma_k}$ can then be written as

$$\sqrt{\Gamma_k} = \sqrt{\sin^2(\theta)\,\Gamma_{\text{ang},11} + \Gamma_{\text{ang},22}} \geq \sqrt{\sin^2(\theta)\,\sigma_\phi^2 + \sigma_\theta^2} = \sigma_k, \tag{S18}$$

where $(\sigma_\phi^2, \sigma_\theta^2)$ are the best-possible variances of measuring $(\phi, \theta)$ calculated from the CRB matrix in angular space; this relationship holds for any molecular orientation $(\phi, \theta, \Omega)$. We compare the angular VUB to the angular CRB averaged over 3D orientation space in Fig. S3. Calculating the average angular VUB $\mathbf{\Gamma}_{\text{ang}}$ is around 32 times faster than calculating the average angular CRB.

### II. EXPERIMENTAL METHODS

Unless stated otherwise, all chemicals were purchased from Sigma-Aldrich and are ACS grade.

#### A. Preparation of amyloid aggregates

The 42 amino-acid residue amyloid-$\beta$ peptide (A$\beta$42) was synthesized and purified by Dr. James I. Elliott (ERI Amyloid Laboratory, Oxford, CT) and dissolved in hexafluoro-2-propanol (HFIP) and sonicated at room temperature for one hour. After flash freezing in liquid nitrogen, HFIP was removed by lyophilization and stored at -20 °C. To further purify the monomeric protein precursors, the lyophilized A$\beta$42 was dissolved in 10 mM NaOH, sonicated for 25 min in a cold water bath and filtered first through a 0.22 µm and then through a 30 kD centrifugal membrane filter (Millipore Sigma, UFC30GV and UFC5030) as described previously [3].

To prepare fibrils, we incubated 10 µM monomeric A$\beta$42 in phosphate-buffered saline (PBS, 150 mM NaCl, 50 mM $Na_3PO_4$, pH 7.4) at 37 °C with 200 rpm agitation for 42-50 hours. The aggregated structures were adsorbed to an ozone-cleaned cell culture chamber (Cellvis, C8-1.5H-N, No. 1.5H, 170 ± 5 µm thickness) for 1 hour immediately after the incubation followed by a rinse using PBS for maximizing adherence of the amyloid to the surfaces of glass-bottom chambers. A PBS solution (200 µL) containing 50 nM Nile Red (Fisher Scientific, AC415711000) was placed into the amyloid-absorbed chambers for transient amyloid binding (TAB) single-molecule orientation localization microscopy (SMOLM).

#### B. Optical instrumentation and imaging procedure

Blinking NR molecules were imaged using a home-built epi-fluorescence microscope equipped with a 100× 1.4 NA oil-immersion objective lens (Olympus, UPlan-SApo 100×). The samples were excited using a 561-nm laser source (Coherent Sapphire, peak intensity at sample ~0.88 kW/cm$^2$) aligned at ~30° tilt from normal for reducing background fluorescence from solution. Fluorescence was collected by the same objective and filtered by a dichroic beamsplitter (Semrock, Di03-R488/561) and a bandpass filter (Semrock, FF01-523/610) followed by separation into two orthogonally-polarized detection channels by a polarizing beamsplitter (Meadowlark Optics, BB-100-VIS). Both channels were captured by a scientific CMOS camera (Hamamatsu, C11440-22CU) with a pixel size of 58.5 × 58.5 nm$^2$ in object space and a conversion gain of 0.49 ADU/photon. Image stacks of 10,000 frames with 20 ms exposure were recorded. A simplified schematic of the imaging system is shown in Fig. S1. Basic photon statistics observed from the NR TAB imaging using the optical system are shown in Fig. S18 and Table S1.

#### C. Background estimation and dual-channel registration

Post-processing on the captured images was performed using custom analysis scripts written in MATLAB (Mathworks, R2019a). The captured raw images were offset corrected by subtracting dark images first. Background photons per pixel were estimated as follows. Each raw image stack was split into sub-stacks and averaged every 200 frames. The averaged images were then separated into two images for considering two polarized channels separately in the following background estimation. We fit each row and column of the images to the sum of two 1D Gaussian functions, with 6 total parameters: 2 sets of amplitudes, centroid positions and standard deviations (widths). We took the average of the two images, i.e., the row-wise and column-wise Gaussian fits, for the following spatial filtering. Broad background fluorescence profiles in each sub-stack and channel were estimated by applying a biorthogonal wavelet filter (level 6 of 'bior6.8' as input parameters to the *wavedec2* and *wdencmp* functions in MATLAB) to the averaged results.

A registration process was also required for analyzing the two orthogonally-polarized fluorescence images simultaneously captured by the camera. The geometric transformation between the two channels on the sCMOS camera was calibrated using fluorescent beads (Thermo Fisher Scientific, FluoSpheres, 0.1 µm, 580/605, F8801) spin-coated on an ozone-cleaned coverslip (Marienfeld, No. 1.5H, 22 × 22 mm, 170 ± 5 µm thickness). Approximately 82,000 - 866,000 photons per bead were detected with 20 ms exposure time. We imaged each bead over 10 frames, and localized bead positions using the ThunderSTORM plugin [4] within ImageJ [5] after subtracting dark images. Calculated bead positions were then averaged across multiple frames for improving localization precision.

Next, all possible lines joining pairs of bead positions across the two channels were drawn. Control point pairs for two-channel registration were selected by comparing the obtained lines, and keeping the largest ensemble of them with similar lengths and slopes. To create the two-channel registration map, coefficients of a global 2D polynomial transformation function were calculated using the control points as input to the *fitgeotrans* function included with MATLAB. To remove small and spatially-varying registration error due to system drift between measurements, we further refined the registration map by re-calculating the global 2D polynomial transformation using single NR molecule positions on the camera localized by ThunderSTORM. Note that the ThunderSTORM localizations of NR molecules were only used to refine the registration map here. Only NR localizations with high localization precision ($< 10$ nm) in each channel were designated as control points.

#### D. Localization and orientation estimation of single molecules

The locations and orientations of NR molecules reported in this paper were estimated simultaneously using a sparsity-promoting maximum likelihood estimator [6, 7]. Briefly, the object space is represented by a rectangular lattice of grid points with spacing equal to the camera pixel size (58.5 nm). Each grid point may contain at most a single molecule parameterized by brightness, position offsets, and six orientational second moments.

To robustly estimate the number of underlying molecules and their parameters in the presence of image overlap, we use a regularized maximum likelihood exploiting a groupsparsity norm to estimate the parameters of each grid point. The algorithm begins by estimating the strength (i.e., brightness) of each of the second moments $\tilde{m}_j$ independently at all object grid points. We next pool together localizations (i.e., their brightnesses and position offsets) across the six second moments to identify the most likely molecules in the object space. Once we identify these molecules, we solve a constrained maximum likelihood to minimize systematic biases induced by the sparsity norm, yielding estimates of the brightnesses, locations, and orientations (second moments $\tilde{\boldsymbol{m}}$ of all molecules in the image). We remove localizations with signal estimates less than 200 photons detected to eliminate unreliable localizations.

The estimated second-moment vectors $\tilde{\boldsymbol{m}}$ were next projected to the first-moment orientation space (azimuthal angle $\phi \in [-\pi, \pi)$, polar angle $\theta \in [0, \pi/2]$, and wobbling area $\Omega \in [0, 2\pi]$ of a transition dipole moment $\boldsymbol{\mu}$) by a weighted least-square estimator:

$$(\phi, \theta, \Omega) = \underset{\phi', \theta', \Omega'}{\arg\min} \left(\tilde{\boldsymbol{m}} - \boldsymbol{m}(\phi', \theta', \Omega')\right)^T \boldsymbol{J} \left(\tilde{\boldsymbol{m}} - \boldsymbol{m}(\phi', \theta', \Omega')\right) \tag{S19}$$

such that

$$\boldsymbol{m}(\phi, \theta, \Omega) = \left[\langle\mu_x^2\rangle, \langle\mu_y^2\rangle, \langle\mu_z^2\rangle, \langle\mu_x\mu_y\rangle, \langle\mu_x\mu_z\rangle, \langle\mu_y\mu_z\rangle\right]^T, \tag{S20}$$

$$\begin{aligned} \langle\mu_x^2\rangle &= \gamma\mu_x^2 + (1-\gamma)/3, \quad \langle\mu_x\mu_y\rangle = \gamma\mu_x\mu_y, \\ \langle\mu_y^2\rangle &= \gamma\mu_y^2 + (1-\gamma)/3, \quad \langle\mu_x\mu_z\rangle = \gamma\mu_x\mu_z, \\ \langle\mu_z^2\rangle &= \gamma\mu_z^2 + (1-\gamma)/3, \quad \langle\mu_y\mu_z\rangle = \gamma\mu_y\mu_z, \end{aligned} \tag{S21}$$

$$[\mu_x, \mu_y, \mu_z] = [\sin\theta\cos\phi, \sin\theta\sin\phi, \cos\theta], \text{ and} \tag{S22}$$

$$\gamma = 1 - \frac{3\Omega}{4\pi} + \frac{\Omega^2}{8\pi^2}, \tag{S23}$$

where $\gamma$ is the rotational constraint [8] and $\boldsymbol{J}$ is the FI matrix calculated from the basis images $\boldsymbol{B}$ as in Eqn. (S4). Here, $\tilde{\boldsymbol{m}}$ and $\boldsymbol{m}$ denote second moment outputs of the maximum likelihood estimator and the weighted least-square estimator respectively. The FI matrix assigns weights to each orientational component $m_j$ inversely proportional to the expected measurement variance $\boldsymbol{R}$ of the polarized PSF. We minimized Eqn. (S19) using the *fmincon* function in MATLAB. If the localization was within a fibril region of interest (Sec. IIII E), then the backbone orientation of the nearest fibril section was used as the initial point of the minimization of Eqn. (S19); otherwise the eigenvector corresponding to the largest eigenvalue of the second moment matrix [9] was assigned as the initial orientation. We have validated and characterized our estimator by using synthetic data and fluorescent beads (Figs. S9 - S14).

#### E. Selection of fibril region of interest and measurement of fibril width

Single-molecule localization microscopy (SMLM) images of amyloid fibrils were generated by binning all single-molecule (SM) localizations within $20 \times 20$ nm$^2$ bins (Fig. S17(a)). Within each field of view, fibrils were identified and segmented into separate regions of interest (ROIs) to compute NR blinking and orientation statistics. Briefly, the SMLM image was converted into a binary image using a localization threshold of 2 localizations/bin and the largest connected structure within a field of view was identified using the *bwconncomp* function in MATLAB. The boundary of the ROI was detected by *bwtraceboundary*, and fibril localizations within the boundary were isolated by *inpolygon* in MATLAB. In order to detect fibril backbones, the SMLM images of isolated fibrils were smoothed by a Gaussian filter and saved as a single tif image. The morphology of each fibril backbone (i.e., a set of points that form a 1D curve in the xy plane) was measured using a ridge detection plugin [10] within ImageJ (Fig. S17(b)). Next, the nearest section of backbone was assigned to each NR localization, which was used for both converting second-moment estimates to first-moment orientations (Sec. IIII D) and comparing NR orientations within the ROI to the fibril superstructure (Figs. 3, S20 - S22).

The full-width at half-maximum (FWHM) of each fibril was measured to characterize their apparent widths (Fig. S19). To measure FWHM, we fit fibril cross-sections, aligned perpendicular locally to each fibril backbone, to 1D Gaussian distributions. Width measurements were performed in intervals of ~17 nm over the entire length of each fibril, thereby building robust statistics for the morphology of the reconstructed fibrils and bundles thereof. Note that the measured apparent widths are wider than the actual widths of the fibrils due to the localization precision of our estimator (Fig. S9).

#### F. Synthetic data

We generated images of molecules via a vectorial image-formation model [11], assuming the orthogonally polarized standard PSF. We used an emission wavelength of 610 nm,

NA = 1.4, a spatially uniform background, and a detector pixel size $58.5 \times 58.5$ nm$^2$ in object space. A mismatched sample refractive index (RI = 1.334) was assumed to mimic the experimental conditions of TAB SMOLM imaging of NR at the coverslip interface within water (Fig. S4).

A molecule is modeled as a dipole rotating uniformly within a cone characterized by a center orientation $(\phi_0, \theta_0)$ and a wobbling area $\Omega_0$. A rotationally fixed dipole corresponds to $\Omega_0 = 0$ sr, while $\Omega_0 = 2\pi$ sr represents an isotropic emitter or a rotationally free molecule (Figs. S9 - S13). Note that our estimator successfully detects all individual emitters if their SBRs are greater than or equal to 300 total signal photons and 2 background photons per pixel.

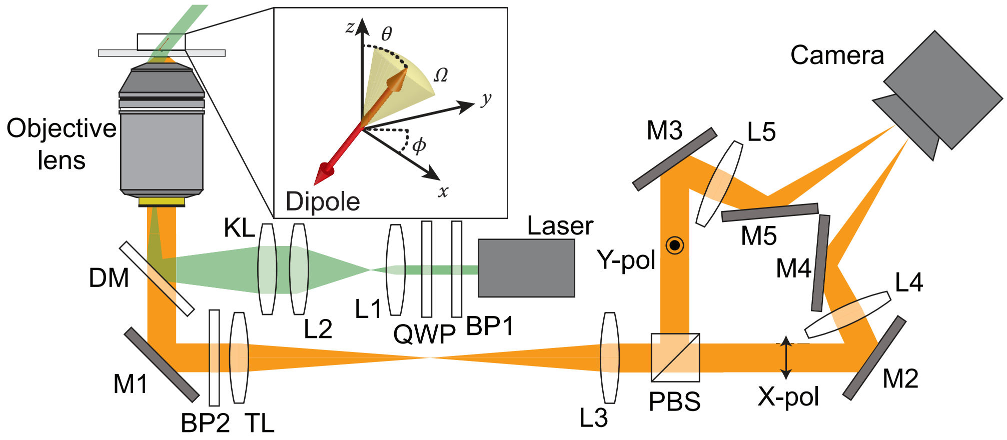

FIG. S1. Experimental setup for SMOLM using the polarized standard PSF. An excitation laser is circularly-polarized and expanded before being coupled into an objective for highly-inclined illumination. Fluorescence from single molecules is collected by the same objective and filtered by a dichroic and a bandpass filter followed by a polarizing beam splitter (PBS) for splitting the light into two orthogonally-polarized channels. After reflection, the two channels (X-pol and Y-pol) are imaged onto different portions of a camera. See Sec. IIII B for details. Inset: The orientation of a transition dipole is parameterized by an azimuthal angle $\phi \in [-\pi, \pi)$, a polar angle $\theta \in [0, \pi/2]$ and a rotational diffusion-associated wobbling area $\Omega \in [0, 2\pi]$. BP1-2, bandpass filters; QWP, quarter wave plate; L1-5, lenses; KL, widefield lens; DM, dichroic mirror; M1-5, mirrors; TL, tube lens.

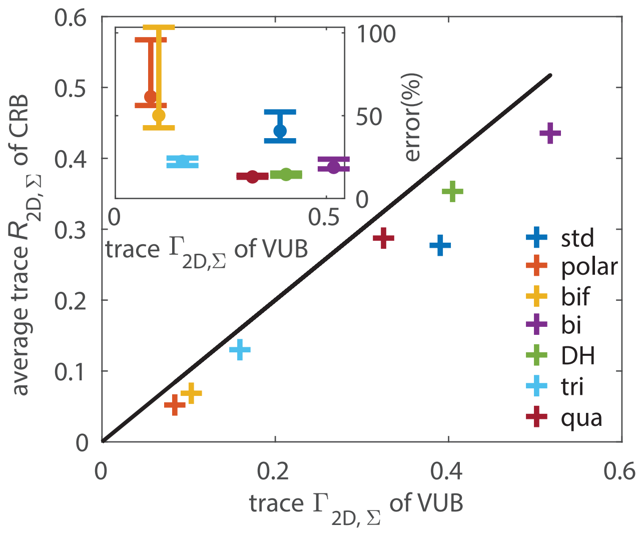

FIG. S2. 2D orientation measurement precision using various PSFs for in-plane oriented molecules at a low signal-to-noise ratio (380 photons detected with 2 background photons/pixel). (a) Average (over the in-plane [xy] orientation space) trace $R_{2D,\Sigma}$ (Eqn. (S13)) of the CRB of the first four orientational second moments $\boldsymbol{m}$ versus the trace $\Gamma_{2D,\Sigma}$ (Eqn. (S14)) of our variance upper bound (VUB) for various PSFs. Blue: unpolarized standard (std), red: polarized standard (polar), yellow: bifocal microscope (bif), purple: bisected (bi), green: double helix (DH), and aqua: tri-spot (tri), maroon: quadrated (qua) PSF. Dark line: $\Gamma_{2D,\Sigma}$ = average $R_{2D,\Sigma}$. Inset: error of $\Gamma_{2D,\Sigma}$ relative to average $R_{2D,\Sigma}$. Bars indicate minimum/maximum error relative to the mean (circle) over all in-plane dipole orientations.

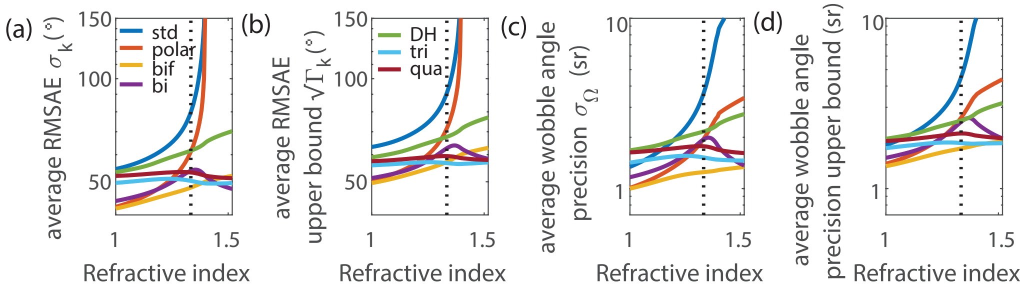

FIG. S3. Precision of measuring the angular first-moment orientation of a molecule at various refractive index interfaces. (a) Average root-mean-square angular error (RMSAE) $\sigma_k$ (Eqn. (S18)), (b) average RMSAE upper bound $\sqrt{\Gamma_k}$ (Eqn. (S18)), (c) average wobble angle precision $\sigma_\Omega$, and (d) average upper bound on wobble angle precision over 3D orientation space for a molecule at various refractive index interfaces. Dotted line: refractive index of water. Blue: unpolarized standard (std), red: polarized standard (polar), yellow: bifocal microscope (bif), purple: bisected (bi), green: double helix (DH), and aqua: tri-spot (tri), maroon: quadrated (qua) PSF.

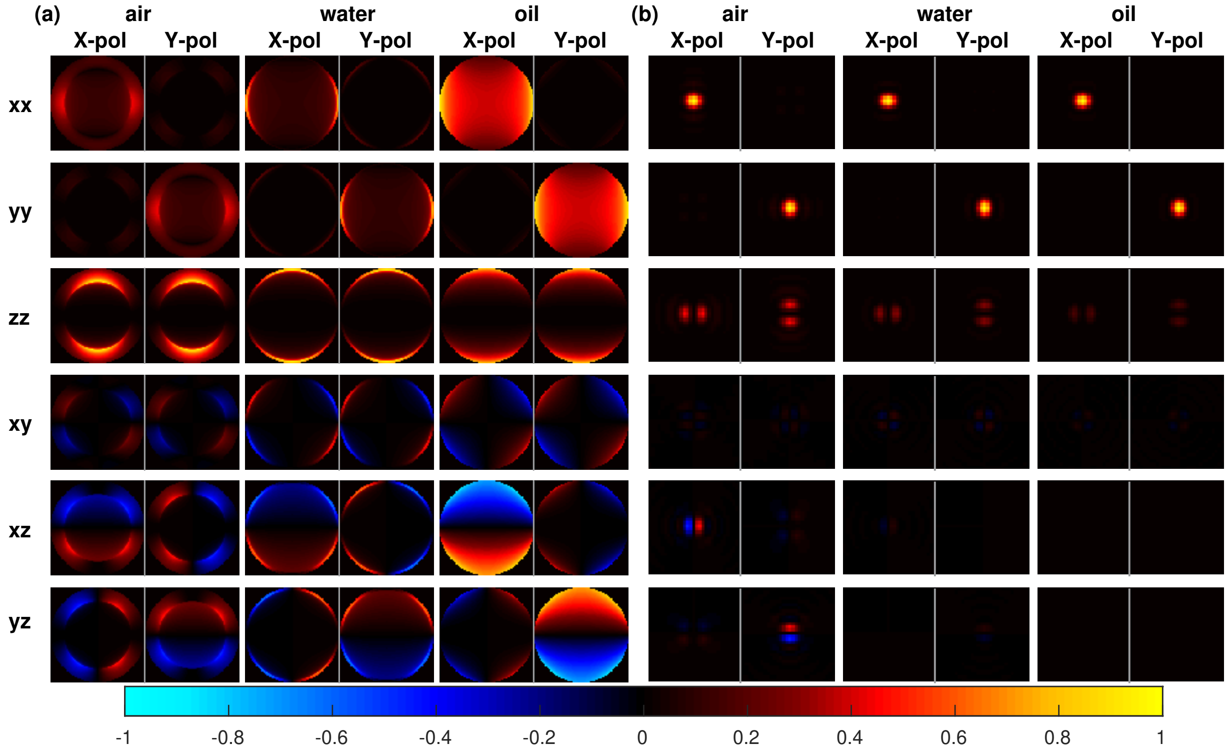

FIG. S4. Simulated polarized orientational basis images (rows, top to bottom: $\boldsymbol{B}_{xx}$, $\boldsymbol{B}_{yy}$, $\boldsymbol{B}_{zz}$, $\boldsymbol{B}_{xy}$, $\boldsymbol{B}_{xz}$, $\boldsymbol{B}_{yz}$) at the (a) back focal plane and (b) image plane of a microscope using a vectorial image-formation model [11] for a molecule at an air-glass, water-glass and oil-glass refractive index interface. Images are normalized in each column.

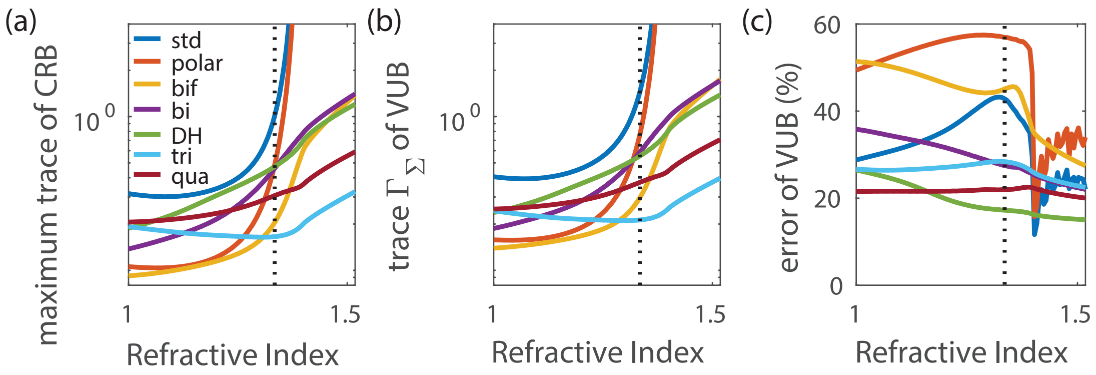

FIG. S5. Comparison of CRB and VUB for emitters at various refractive index interfaces. (a) Maximum trace of CRB matrix $\boldsymbol{R}$, (b) trace $\Gamma_\Sigma$ of VUB and (c) error of VUB relative to the maximum trace of CRB for measuring the orientation of a molecule at various refractive index interfaces. Dotted line: refractive index of water. Blue: unpolarized standard (std), red: polarized standard (polar), yellow: bifocal microscope (bif), purple: bisected (bi), green: double helix (DH), and aqua: tri-spot (tri), maroon: quadrated (qua) PSF.

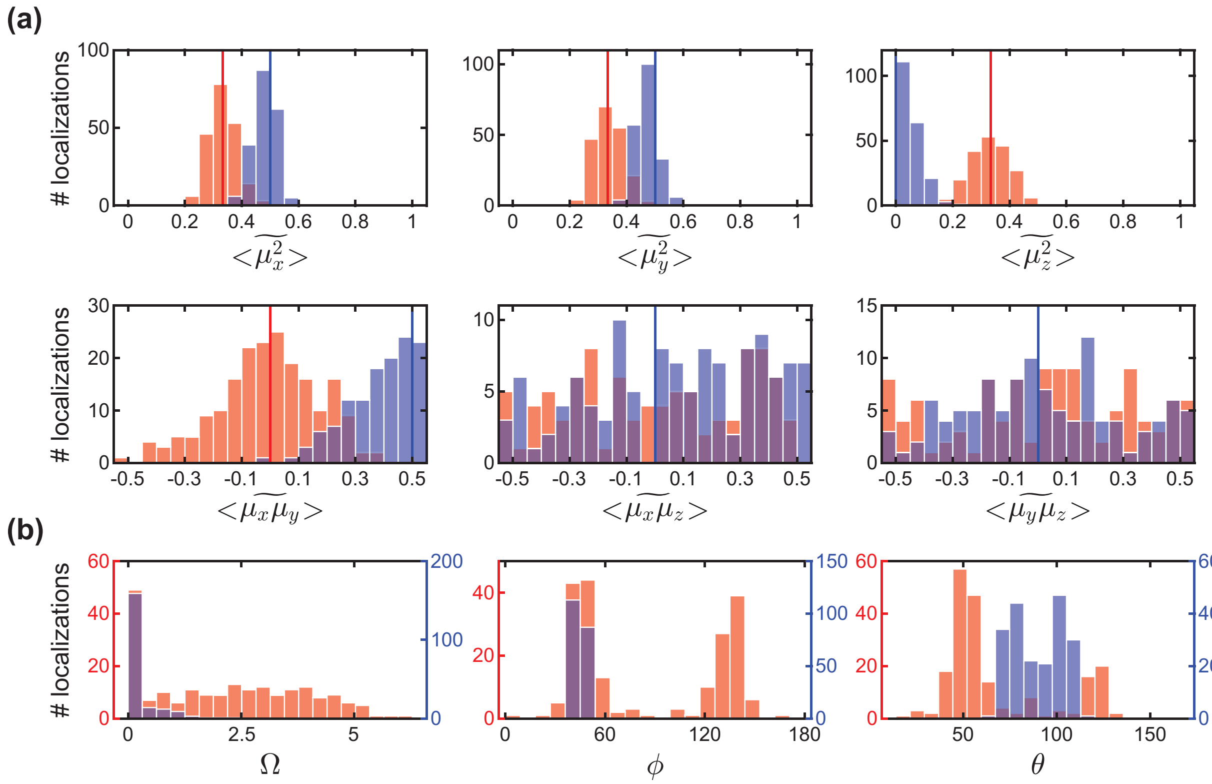

FIG. S6. Resolvability of a fixed molecule versus an isotropic emitter using our maximum likelihood estimator followed by a weighted least-square estimator. (a) Estimated second moments $\tilde{m}$ using the polarized standard PSF and the maximum likelihood estimator described in Sec. IIII D for a fixed molecule with orientation (blue, $\Omega_0 = 0$ sr, $\theta_0 = 90°$, $\phi_0 = 45°$, Fig. 1(d)) and an isotropic emitter (red, $\Omega_0 = 2\pi$ sr, Fig. 1(e)). At each orientation, 200 independent images were generated as described in Sec. IIII F with $s_0 = 380$ signal photons, $b_0 = 2$ background photons/pixel. The vertical lines depict the ground truth second moments ($\boldsymbol{m}_0$) for the fixed molecule (blue) and the isotropic emitter (red). The estimator allows the cross terms ($\langle \widetilde{\mu_x\mu_y} \rangle, \langle \widetilde{\mu_x\mu_z} \rangle, \langle \widetilde{\mu_y\mu_z} \rangle$) to have non-physical values (e.g., $\langle \widetilde{\mu_x\mu_y} \rangle < -0.5$ or $\langle \widetilde{\mu_x\mu_y} \rangle > 0.5$). Note that the estimator gives unbiased and precise measurements of the first four second moments but wide distributions for the last two moments. This behavior is consistent with the discussion in the main text and Fig. 1; the polarized standard PSF contains little sensitivity to $\langle \widetilde{\mu_x\mu_z} \rangle$ and $\langle \widetilde{\mu_y\mu_z} \rangle$. (b) Projected mean orientation ($\phi$,$\theta$) and wobbling area ($\Omega$) of each estimate obtained from $\tilde{m}$ using the weighted least-square estimator described in Sec. IIII D for the fixed in-plane molecule (blue) and the isotropic emitter (red). Although $\Omega$ estimates of the isotropic emitter show unavoidable bias and low precision due to photon shot noise (Fig. S13(a)) [12], the estimates demonstrate that the fixed molecule and isotropic emitter are easily resolved by our method. Note that measurements of $\theta$ and $\phi$ have little significance for the isotropic emitter (red).

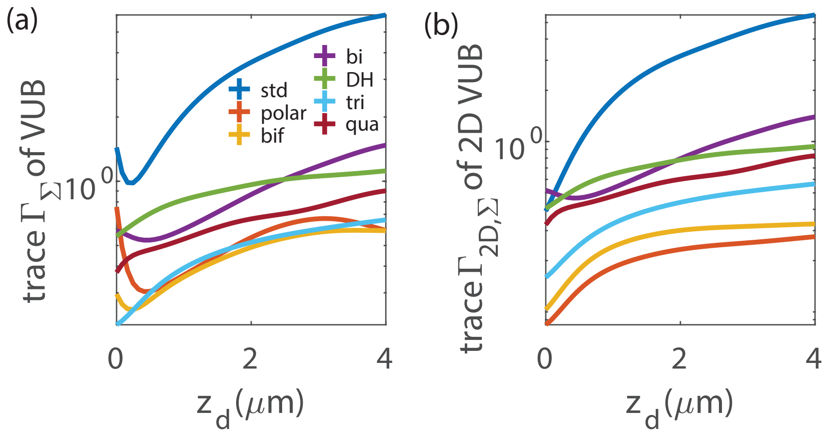

FIG. S7. Orientation measurement precision for emitters at various depths within water. (a) Trace $\Gamma_\Sigma$ of VUB and (b) trace $\Gamma_{2D,\Sigma}$ (Eqn. (S14)) of 2D VUB for emitters at various depths $z_d$ within water, relative to the coverlip at $z_d = 0$. At each emitter depth $z_d$, each PSF was refocused to achieve best focus. Blue: unpolarized standard (std), red: polarized standard (polar), yellow: bifocal (bif), purple: bisected (bi), green: double helix (DH), and aqua: tri-spot (tri), maroon: quadrated (qua) PSF.

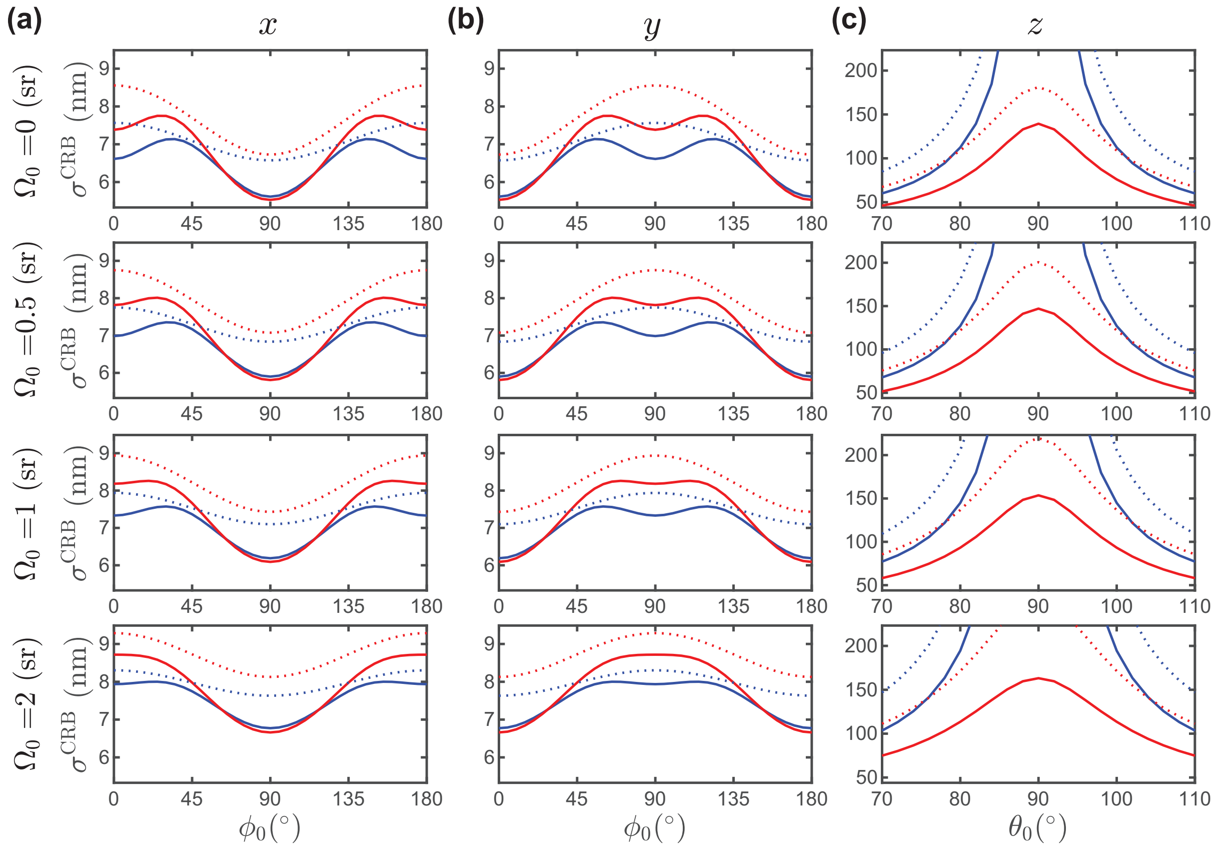

FIG. S8. Cramér–Rao bounds for estimating the 3D position along (a) $x$, (b) $y$, and (c) $z$ of a single molecule using the polarized (solid) and unpolarized (dashed) standard PSFs. The bounds are computed for a focused molecule ($z = 0$ nm) at various orientations ($\Omega_0 = 0 - 2$ sr , $\phi_0 = 0 - 180^\circ$, $\theta_0 = 70 - 110^\circ$) within a matched medium (RI = 1.518, blue) and at a mismatched refractive index interface (RI = 1.334, red). Calculations were performed for $s_0 = 380$ signal photons and $b_0 = 2$ background photons/pixel. Note that the square root of the CRB for $x$ and $y$ estimations were averaged over all simulated $\theta_0$ since the estimation precisions are mostly constant for the range of $\theta_0$ examined here. Similarly, the square root of the CRB for $z$ estimation was averaged over all $\phi_0$.

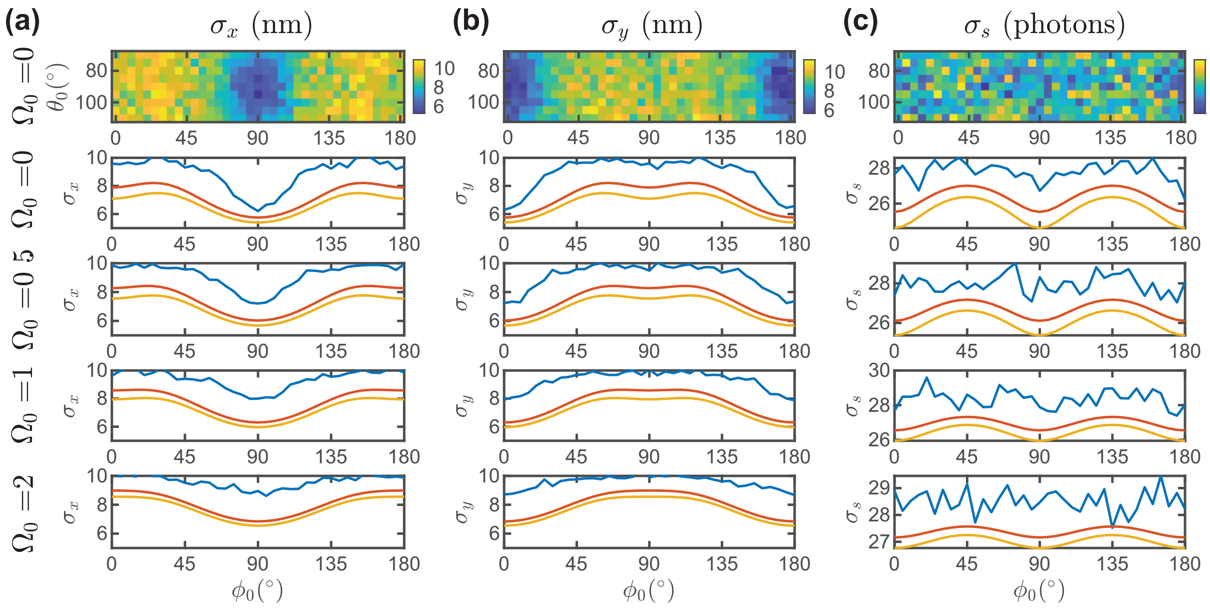

FIG. S9. SMOLM measurement precision for 2D position (a) $\sigma_x$ and (b) $\sigma_y$ and (c) brightness $\sigma_s$ determined by repeatedly localizing dipoles at various orientations ($\Omega_0 = 0 - 2$ sr, $\phi_0 = 0 - 180°$, $\theta_0 = 70 - 110°$) in synthesized polarized standard PSF images. At each orientation, 200 independent images were generated as described in Sec. IIII F with $s_0 = 380$ photons, $b_0 = 2$ background photons/pixel (a total of 266,400 images across 1,332 different orientations). Positions and photons of simulated molecules were estimated as described in Sec. IIII D, and the measurement standard deviation was computed for each ground-truth orientation. Note that the estimator detects all emitters at this SBR, since the localization and signal estimation precisions have little correlation with $\theta_0$, estimation performance was accumulated over all simulated $\theta_0$ and compared against $\phi_0$ and $\Omega_0$ (blue lines in the bottom 4 rows). The square root of the CRB for $x$, $y$, and $s$ estimations are shown for (red) 20° out-of-plane ($\theta_0 = 70°, 110°$) and (orange) in-plane ($\theta_0 = 90°$) orientations.

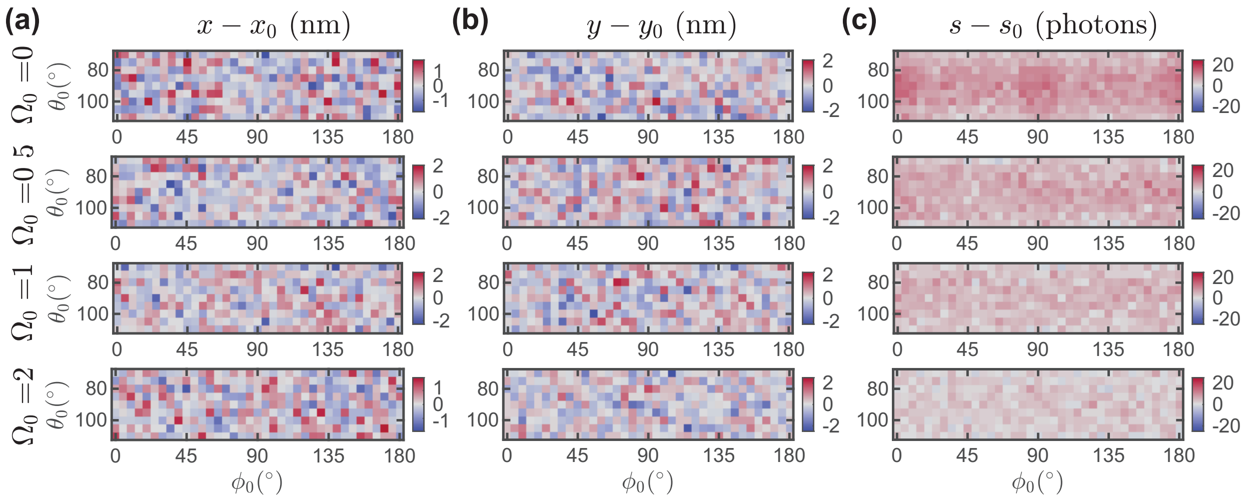

FIG. S10. SMOLM measurement bias for 2D position (a) $x - x_0$ and (b) $y - y_0$ and (c) brightness $s - s_0$ determined by repeatedly localizing dipoles at various orientations ($\Omega_0 = 0 - 2$ sr, $\phi_0 = 0 - 180°$, $\theta_0 = 70 - 110°$) in synthesized polarized standard PSF images. At each orientation, 200 independent images were generated as Sec. IIII F with $s_0 = 380$ photons, $b_0 = 2$ background photons/pixel (a total of 266,400 images across 1,332 different orientations). Positions and photons of simulated molecules were estimated as described in Sec. IIII D, and the measurement bias was computed by averaging the measurement errors at each ground-truth orientation.

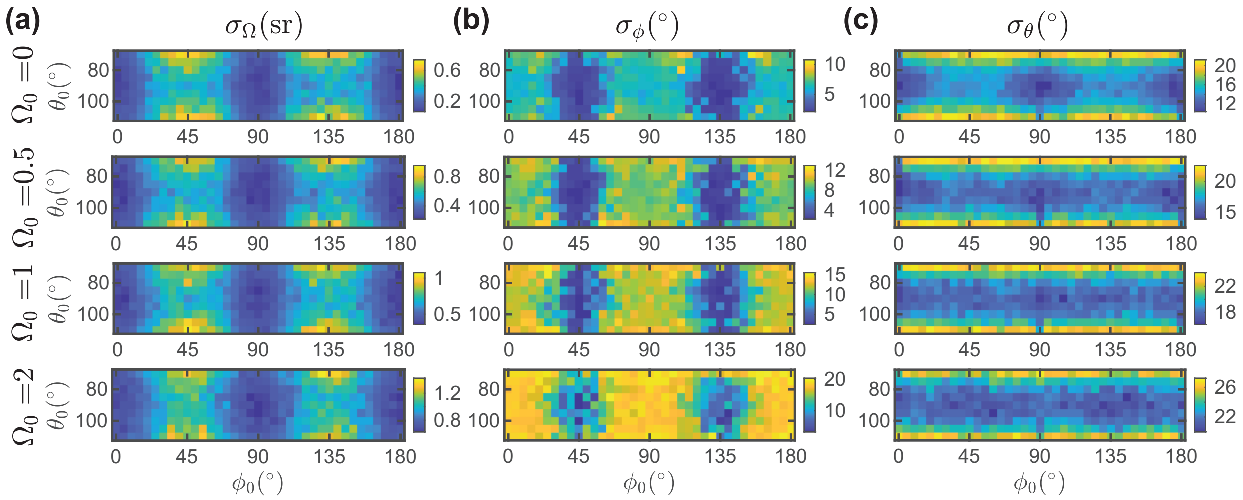

FIG. S11. SMOLM measurement precision for (a) wobbling area $\sigma_\Omega$, (b) azimuthal orientation $\sigma_\phi$, and (c) polar orientation $\sigma_\theta$ determined by repeatedly localizing dipoles at various orientations ($\Omega_0 = 0 - 2$ sr, $\phi_0 = 0 - 180°$, $\theta_0 = 70 - 110°$) in synthesized polarized standard PSF images. At each orientation, 200 independent images were generated as Sec. IIII F with $s_0 = 380$ photons, $b_0 = 2$ background photons/pixel (a total of 266,400 images across 1,332 different orientations). Orientations of simulated molecules were estimated as described in Sec. IIII D, and the measurement standard deviation was computed at each ground-truth orientation.

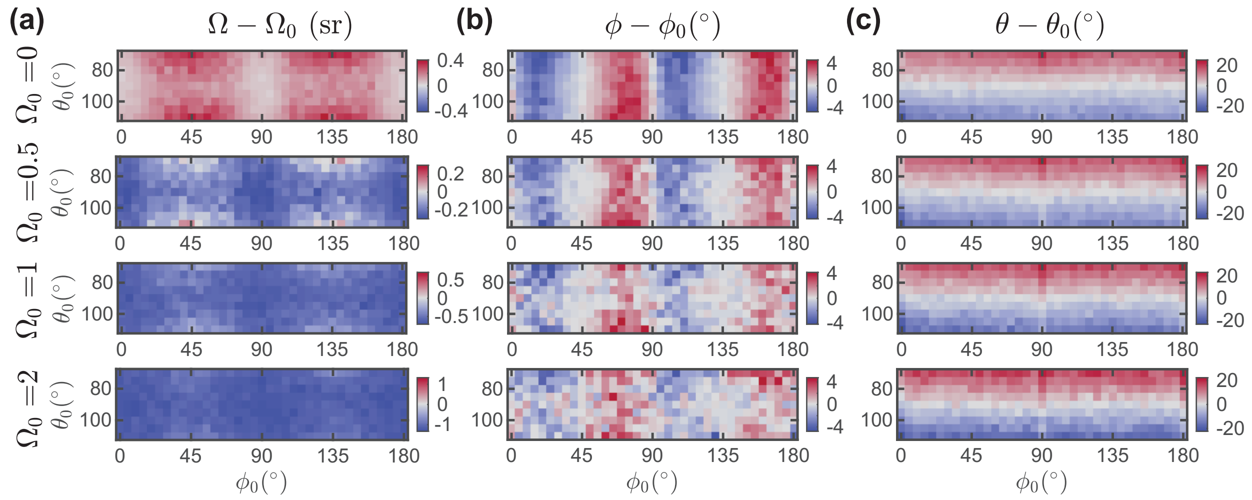

FIG. S12. SMOLM measurement bias for (a) wobbling area $\Omega - \Omega_0$, (b) azimuthal orientation $\phi - \phi_0$, and (c) polar orientation $\theta - \theta_0$ determined by repeatedly localizing dipoles at various orientations ($\Omega_0 = 0 - 2$ sr, $\phi_0 = 0 - 180°$, $\theta_0 = 70 - 110°$) in synthesized polarized standard PSF images. At each orientation, 200 independent images were generated as Sec. IIII F with $s_0 =$ 380 photons, $b_0 = 2$ background photons/pixel (a total of 266,400 images across 1,332 different orientations). Orientations of simulated molecules were estimated as described in Sec. IIII D, and the measurement bias was computed by averaging the measurement errors at each ground-truth orientation.

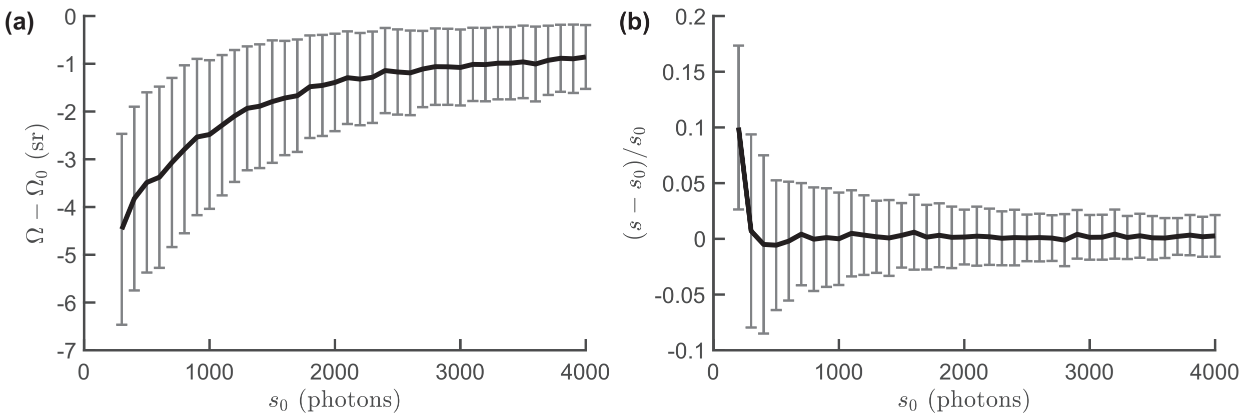

FIG. S13. SMOLM measurement bias for (a) wobbling area $\Omega - \Omega_0$ and (b) brightness $s - s_0$ determined by repeatedly localizing isotropic emitters ($\Omega_0 = 2\pi$ sr) with various brightnesses ($s_0 = 200 - 4000$ photons) in synthesized images. At each brightness, 5000 independent images were generated as Sec. IIII F with $b_0 = 2$ background photons/pixel (a total of 195,000 images). The wobbling areas and brightnesses of simulated molecules were estimated as described in Sec. IIII D, and the estimation bias was computed by averaging the measurement errors at each brightness condition $s_0$. Error bars represent standard deviations of the estimates. Systematic underestimation of the wobbling area, i.e., apparent rotational diffusion that is smaller than the ground truth, is attributed to Poisson shot noise and is consistent with theoretical models [12]. Overestimation of the brightness at 200 photons is due to the filtering (removal) of weak localizations with less than 200 photons detected.

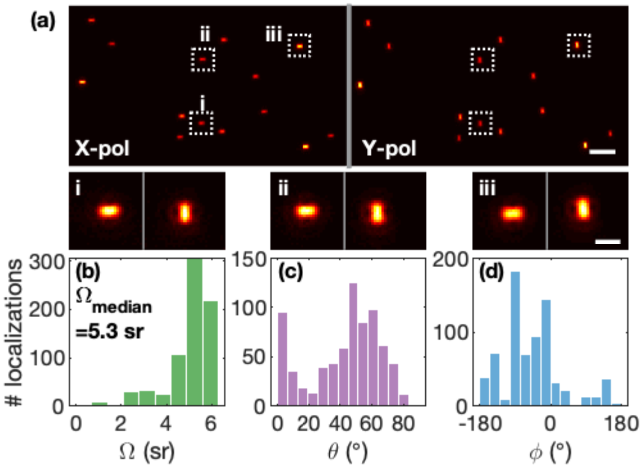

FIG. S14. (a) Raw images of fluorescent beads (Thermo Fisher Scientific, FluoSpheres, 0.1 μm, 580/605, F8801). Scale bar: 2 μm. Inset: zoomed x-y polarization channel images of three beads. Scale bar: 400 nm. Measured (b) $\Omega$, (c) $\theta$, and (d) $\phi$ of fluorescent beads (730 localizations of 73 beads) with an average of 300,000 photons detected and 3 background photons/pixel. Our estimation results indicate that the fluorescent beads emit light mostly isotropically ($\Omega \approx 2\pi$ sr), which is consistent with our expectation on emitters with a large number of fluorophores. Note that measurements of $\theta$ and $\phi$ have little significance with such large measured values of $\Omega$.

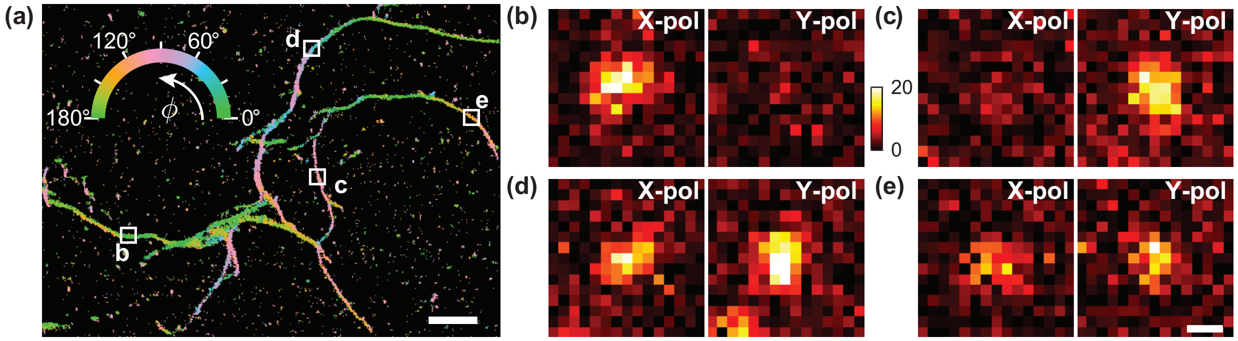

FIG. S15. Fluorescence images of NR molecules on fibrils. (a) TAB SMOLM image, color-coded according to the mean azimuthal ($\phi$) orientation of the NR molecules measured within each bin ($20\times20$ nm$^2$). (b-e) Representative fluorescence raw images of single NR molecules on the fibril network shown in Fig. 2. The observed polarized images of NR molecules exhibit substantial differences depending on the orientation of the underlying fibril structures [(b) $\phi_{\text{avg}} = 176°$, (c) $\phi_{\text{avg}} = 108°$, (d) $\phi_{\text{avg}} = 50°$, and (e) $\phi_{\text{avg}} = 140°$]. Color bar: photons/pixel. Images contain (b) 290, (c) 334, (d) 523, and (e) 336 detected photons across both polarized images. Scale bars: (a) 1 µm, (e) 200 nm.

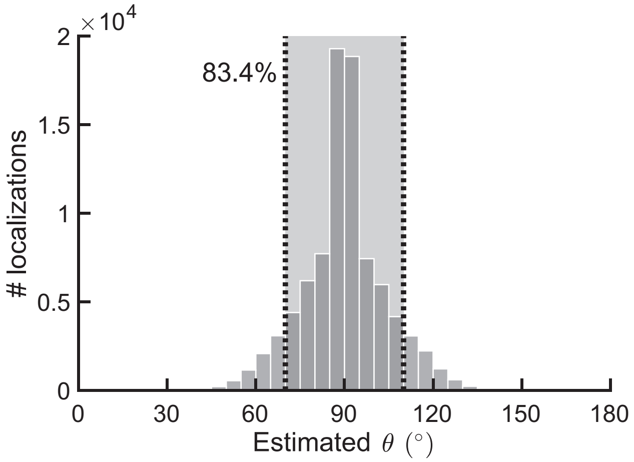

FIG. S16. The estimated polar orientations $\theta$ of Nile red molecules on fibrils show small out-of-plane angles. NR molecules with near in-plane polar orientations ($70° < \theta < 110°$) dominate ($> 83\%$) the distribution. These data correspond to the fibrils shown in Fig. 2.

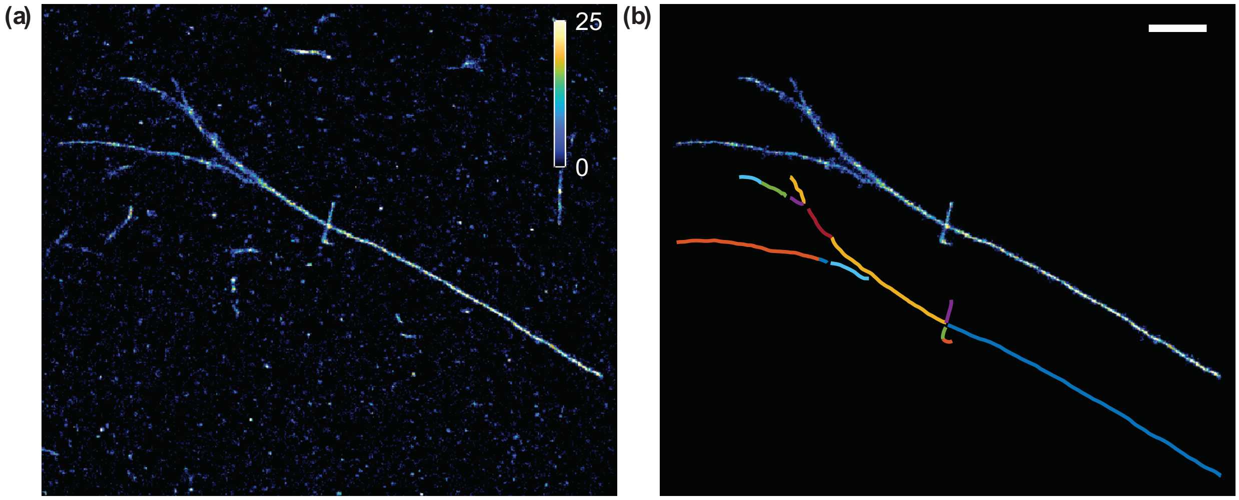

FIG. S17. Fibril region of interest (ROI) and backbone ridge detection. (a) SMLM image of the fibril bundles shown in Figs. 3(a-d). (b) Isolated fibril ROI and the corresponding detected ridges (i.e., fibril backbones) based on the ROI selection method described in Sec. IIII E. The ridges are individually indexed by color, but this indexing (classification) was ignored in our analyses. Scale bar: 1 µm. Color bar: localizations/bin.

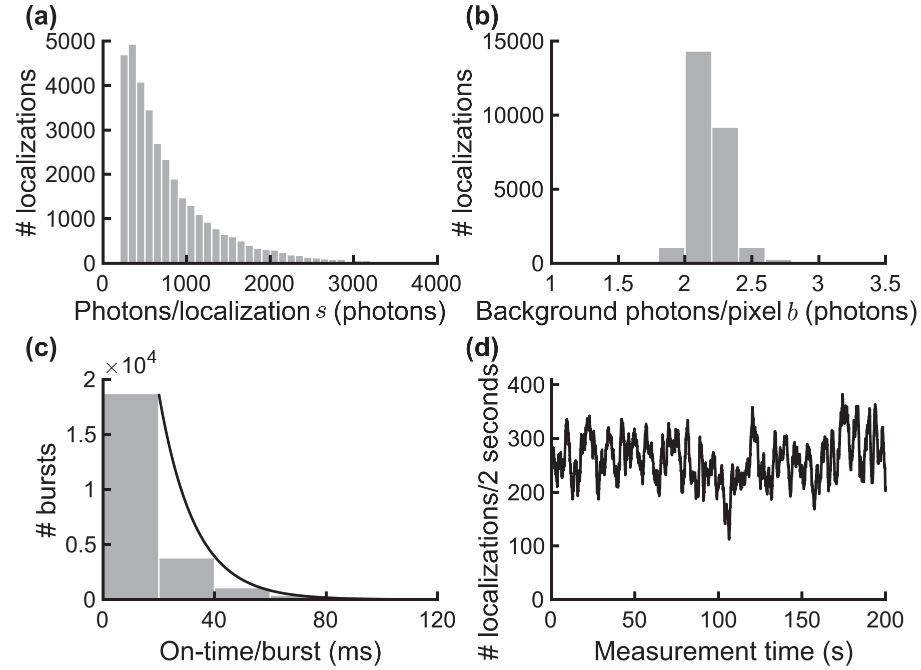

FIG. S18. Photophysics of Nile red localization and blinking events. (a) Photons detected per localization $s$, (b) background photons per pixel $b$ and (c) on-time of NR bursts on amyloid fibrils. The black solid line in (c) depicts a fit to an exponential decay. The median numbers of photons detected per localization and background photons per pixel are 600 and 2.2, respectively; the time constant of the exponential fit ($\tau_{\text{on}}$) is 13 ms (Table S1). (d) Localization rate per 2 seconds of NR on amyloid fibrils. Continuous replenishment of NR molecules from the imaging buffer to amyloid surfaces prevents obvious photobleaching and image degradation over the measurement time. These data correspond to the fibril bundles shown in Figs. 3(a-d) in the main text.

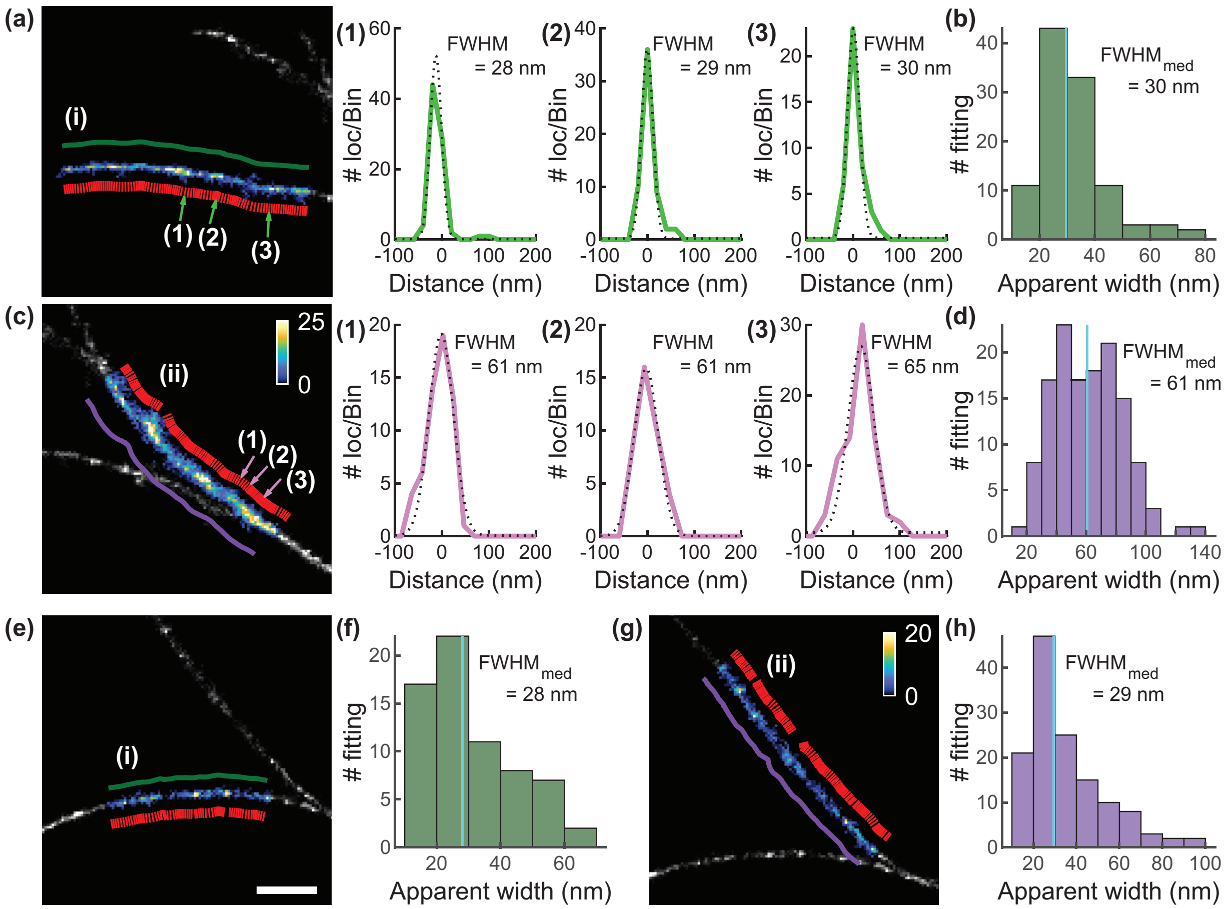

FIG. S19. Measurements of fibril apparent widths for the regions in Fig. 3. (a) Zoomed-in SMLM image of the fibril shown in Figs. 3(a-d)(i). The green line depicts the detected backbone in this region. The red lines represent each cross-sectional profile aligned perpendicular to the ridge. Three representative width profiles (green) and their Gaussian fits (dotted black) are shown in (1-3) with the estimated FWHM. (b) All apparent fibril widths within the region shown in (a). The median FWHM is 30 nm. (c) Zoomed-in SMLM image of the fibril bundle shown in Figs. 3(a-d)(ii) and its representative cross-sectional profiles. (d) All apparent fibril widths within the region shown in (c). The median FWHM is 61 nm. (e-h) Zoomed-in SMLM images of the fibrils shown in Figs. 3(e-h)(i) and (ii), and the distribution of fitted FWHMs in the two regions. The median apparent widths are (i) 28 nm and (ii) 29 nm respectively. Scale bar: 500 nm. Color bars: localizations/bin. Orientation-localization data are available in Dataset 1 [13].

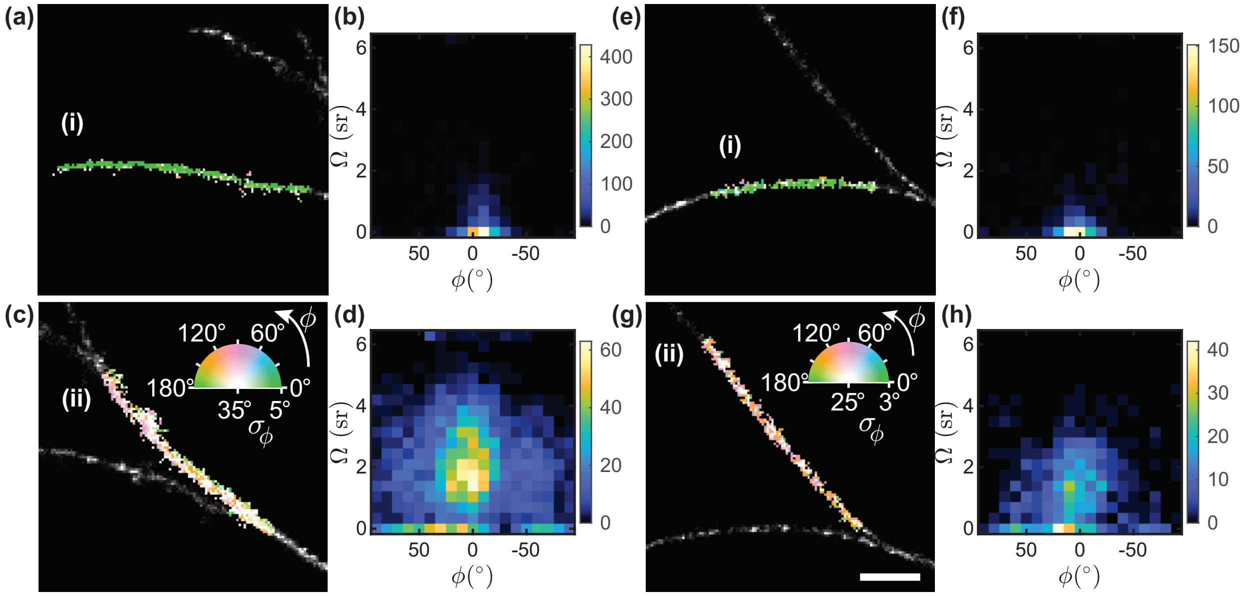

FIG. S20. Correlation between estimates of azimuthal orientation $\phi$ and wobbling area $\Omega$. (a,c) and (e,g) Zoomed-in TAB SMOLM images corresponding to Figs. 3(a-d) and 3(e-h) respectively, color-coded according to the mean azimuthal angle ($\phi$) and standard deviation of the orientations ($\sigma_\phi$) measured within each bin ($20 \times 20$ nm$^2$). Scale bar: 500 nm. (b,d) and (f,h) 2D histograms of the estimated $\Omega$ vs $\phi$ within the regions depicted by (a,c) and (e,g). Color bar: localizations per orientation. We observe broader distributions of $\phi$ and $\Omega$ in (d) and (h) than those in (b) and (f), implying that NR orientations in (ii) are more heterogeneous comparing to those in (i). Orientation-localization data are available in Dataset 1 [13].

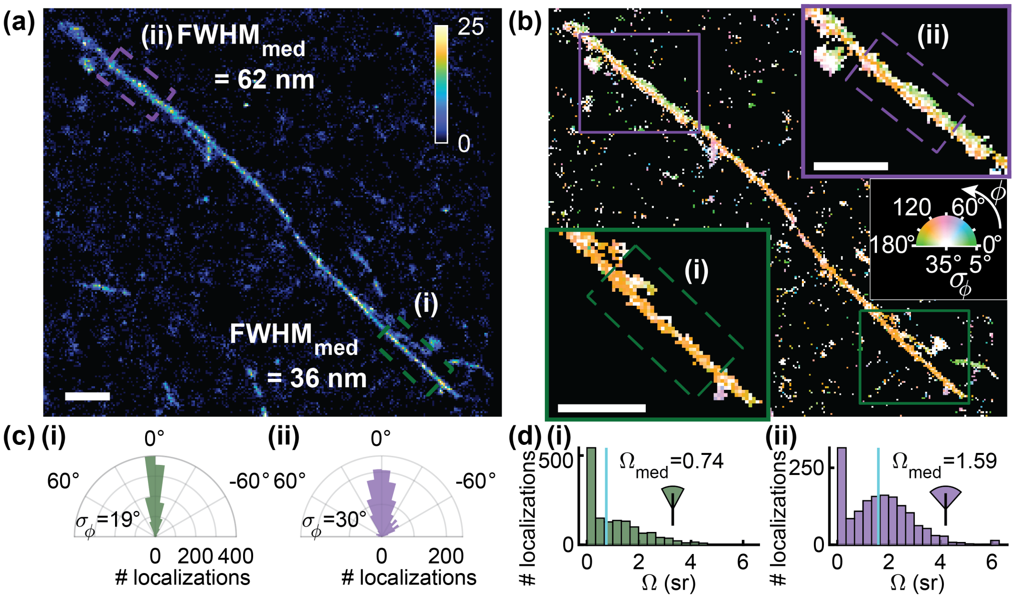

FIG. S21. Structural heterogeneity of A$\beta$42 fibrils revealed by TAB SMOLM imaging I. (a) SMLM image of a fibril. Color bar: localizations per bin ($20 \times 20$ nm$^2$). Apparent fibril widths represented by FWHM are (i) 36 nm and (ii) 62 nm in the regions denoted by (i) green and (ii) purple boxes. (b) TAB SMOLM image corresponding to (a), color-coded according to the mean azimuthal angle ($\phi$) and standard deviation of the orientations ($\sigma_\phi$) measured within each bin. Insets: zoomed (i) thin and (ii) thick fibril regions isolated from background structures. (c) Distributions of NR azimuthal orientations relative to the fibril backbone within the regions denoted in (b). The standard deviations ($\sigma_\phi$) in relative backbone angle are (i) 19° and (ii) 30°. (d) Histograms of the measured wobbling area ($\Omega$) corresponding to the localizations in (c). The median wobbling areas ($\Omega_{\text{med}}$) are (i) 0.74 sr and (ii) 1.59 sr. Scale bars: 500 nm. Orientation-localization data are available in Dataset 1 [13].

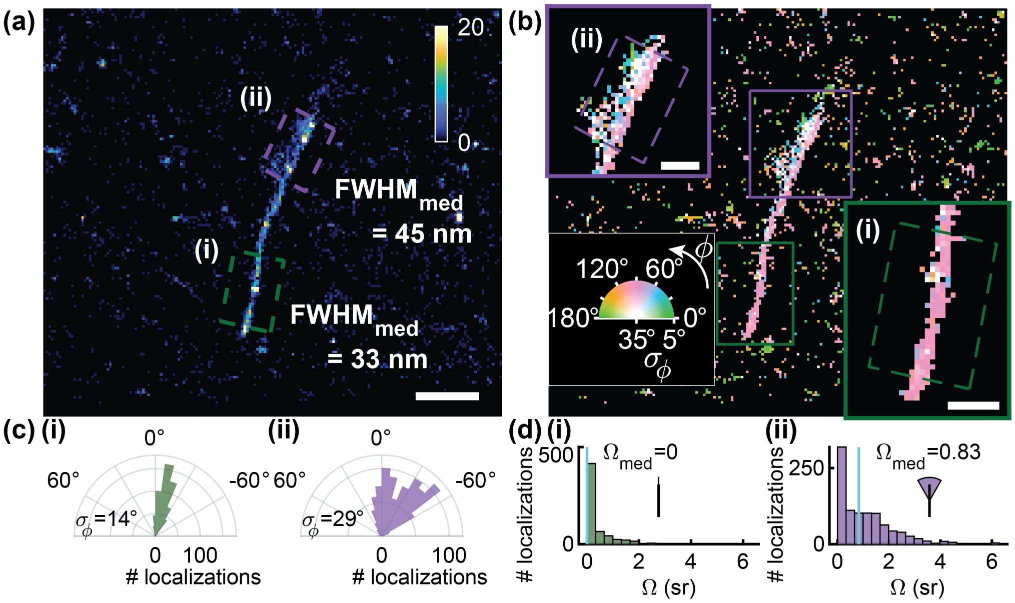

FIG. S22. Structural heterogeneity of A$\beta$42 fibrils revealed by TAB SMOLM imaging II. (a) SMLM image of a fibril. Color bar: localizations per bin ($20 \times 20$ nm$^2$). Scale bar: 500 nm. Apparent fibril widths represented by FWHM are (i) 33 nm and (ii) 45 nm in the regions denoted by (i) green and (ii) purple boxes. (b) TAB SMOLM image corresponding to (a), color-coded according to the mean azimuthal angle ($\phi$) and standard deviation of the orientations ($\sigma_\phi$) measured within each bin. Inset: zoomed (i) thin and (ii) thick fibril regions isolated from background structures. Scale bars: 200 nm. (c) Distributions of NR azimuthal orientations relative to the fibril backbone within the regions denoted in (b). The standard deviations ($\sigma_\phi$) in relative backbone angle are (i) 18° and (ii) 29°. (d) Histograms of the measured wobbling area ($\Omega$) corresponding to the localizations in (c). The median wobbling areas ($\Omega_{med}$) are (i) 0 sr and (ii) 0.83 sr. Orientation-localization data are available in Dataset 1 [13].

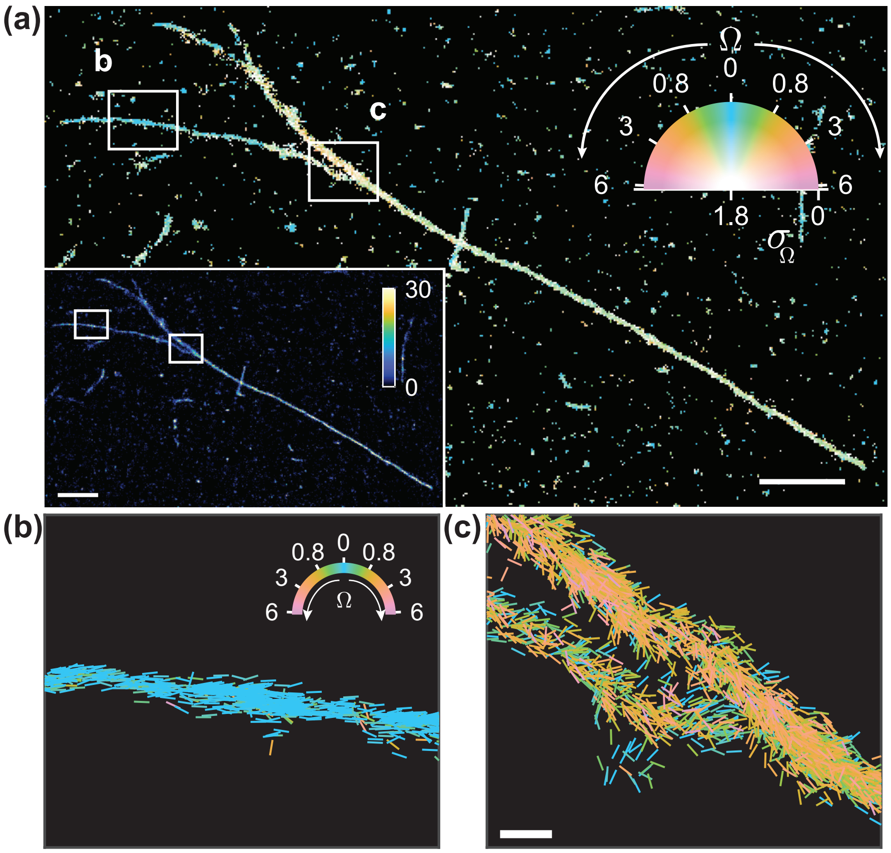

FIG. S23. Estimated wobbling area Ω of NR molecules on amyloid fibrils. (a) TAB SMOLM image, color-coded according to the mean wobbling area (Ω, sr) and standard deviation of the rotational diffusion ($\sigma_\Omega$) measured within each bin. Inset: SMLM image of the fibril bundles. Color bar: localizations per bin ($20 \times 20$ nm$^2$). Scale bar: 1 µm. (b,c) All individual wobbling measurements localized along fibril backbones within the white boxes in (a). The lines represent the direction of the estimated $\phi$ of each localization and are color-coded according to the estimated Ω. Scale bar: 100 nm. These data correspond to the fibril bundles shown in Figs. 3(a-d) in the main text. Orientation-localization data are available in Dataset 1 [13].

TABLE S1. Localization numbers, localization density, photons detected per localization, background photons per pixel, and burst on-time of Nile Red molecules on amyloid structures

| | # Localizations | Localizations/μm (per fibril length) | Photons/loc[1] $s$ | Background photons /pixel[1] $b$ | On-time (ms) $\tau_{\mathrm{on}}$ |
|---|---|---|---|---|---|
| FIG. 2 | 88748 | 2373 | 481 | 2.4 | 14 |
| FIG. 3a-d | 34352 | 2368 | 600 | 2.2 | 13 |
| FIG. 3e-h | 13351 | 1477 | 460 | 1.4 | 17 |
| FIG. S21 | 17425 | 2607 | 566 | 2.5 | 14 |
| FIG. S22 | 3466 | 1833 | 443 | 2.0 | 13 |

[1] Median of each statistic.

[2] Orientation-localization data are available in Dataset 1 [13].

Movie S1 Raw images of single NR molecules transiently binding to A$\beta$42 fibrils. Raw images of single NR molecules transiently binding to A$\beta$42 fibrils. Hot color scale: photons detected per pixel (58.5 $\times$ 58.5 nm$^2$). Corresponding SMLM images of the fibrils are shown to the right by accumulating localizations within 20 $\times$ 20 nm$^2$ bins over time. Color scale: localizations per bin. Scale bar: 1 μm.

Movie S2 Individual azimuthal orientation measurements localized along fibril backbones. (top) TAB SMOLM image of the fibrils shown in Figs. 3(a-d), color-coded according to the mean azimuthal angle ($\phi$) and standard deviation of the orientations ($\sigma_\phi$) measured within each bin. Scale bar: 1 μm. Inset: SMLM image of the corresponding A$\beta$42 fibrils. Color bar: localizations per bin (20 $\times$ 20 nm$^2$). Scale bar: 1 μm. (bottom) All individual orientation measurements localized along the fibril backbones within the white boxes. The lines depict the direction of the estimated $\phi$ angle and are color-coded accordingly. Scale bar: 100 nm.

[1] R. Bhatia, Variational principles for eigenvalues, in *Matrix Analysis* (Springer New York, New York, NY, 1997) pp. 57–83.

[2] A. Nehorai and E. Paldi, Vector-sensor array processing for electromagnetic source localization, IEEE transactions on signal processing **42**, 376 (1994).

[3] K. Spehar, T. Ding, Y. Sun, N. Kedia, J. Lu, G. R. Nahass, M. D. Lew, and J. Bieschke, Super-resolution Imaging of Amyloid Structures over Extended Times by Using Transient Binding of Single Thioflavin T Molecules, ChemBioChem **19**, 1944 (2018).

[4] M. Ovesný, P. Křížek, J. Borkovec, Z. Švindrych, and G. M. Hagen, ThunderSTORM: A comprehensive ImageJ plug-in for PALM and STORM data analysis and super-resolution imaging, Bioinformatics **30**, 2389 (2014).

[5] C. A. Schneider, W. S. Rasband, and K. W. Eliceiri, NIH Image to ImageJ: 25 years of image analysis, Nature methods **9**, 671 (2012).

[6] H. Mazidi, E. S. King, O. Zhang, A. Nehorai, and M. D. Lew, Dense Super-Resolution Imaging of Molecular Orientation Via Joint Sparse Basis Deconvolution and Spatial Pooling, in *IEEE 16th International Symposium on Biomedical Imaging (ISBI)*, 325–329 (2019).

[7] H. Mazidi, T. Ding, and M. D. Lew, `https://github.com/Lew-Lab/RoSE-O`.

[8] O. Zhang, J. Lu, T. Ding, and M. D. Lew, Imaging the three-dimensional orientation and rotational mobility of fluorescent emitters using the tri-spot point spread function, Applied physics letters **113**, 031103 (2018).

[9] A. S. Backer and W. Moerner, Determining the rotational mobility of a single molecule from a single image: a numerical study, Optics express **23**, 4255 (2015).

[10] C. Steger, An unbiased detector of curvilinear structures, IEEE Transactions on Pattern Analysis and Machine Intelligence **20**, 113 (1998).

[11] A. S. Backer and W. Moerner, Extending single-molecule microscopy using optical fourier processing, The Journal of Physical Chemistry B **118**, 8313 (2014).

[12] O. Zhang and M. D. Lew, Fundamental Limits on Measuring the Rotational Constraint of Single Molecules Using Fluorescence Microscopy, Physical Review Letters **122**, 198301 (2019).

[13] M. D. Lew, T. Ding, and T. Wu, `https://osf.io/pe3qu/?view_only=081206495472426889c1055f21971e9a`.
